# Enhancement of trans-cleavage activity of Cas12a with engineered crRNA enables amplified nucleic acid detection

**DOI:** 10.1101/2020.04.13.036079

**Authors:** Long T. Nguyen, Brianna M. Smith, Piyush K. Jain

**Affiliations:** Department of Chemical Engineering, University of Florida, 1006 Center Drive, Gainesville, FL 32611, USA; UF Health Cancer Center, University of Florida, 2033 Mowry Rd., CGRC 463, Gainesville, FL 32608, USA

## Abstract

The CRISPR/Cas12a RNA-guided complexes have a tremendous potential for nucleic acid detection due to its ability to indiscriminately cleave ssDNA once bound to a target DNA. However, the current CRISPR/Cas12a systems are limited to detecting DNA in a picomolar detection limit without an amplification step. Here, we developed a platform with engineered crRNAs and optimized conditions that enabled us to detect DNA, DNA/RNA heteroduplex and methylated DNA with higher sensitivity, achieving a limit of detection of in femtomolar range without any target pre-amplification step. By extending the 3’- or 5’-ends of the crRNA with different lengths of ssDNA, ssRNA, and phosphorothioate ssDNA, we discovered a new self-catalytic behavior and an augmented rate of LbCas12a-mediated collateral cleavage activity as high as 3.5-fold compared to the wild-type crRNA. We applied this sensitive system to detect as low as 25 fM dsDNA from the PCA3 gene, an overexpressed biomarker in prostate cancer patients, in simulated urine over 6 hours. The same platform was used to detect as low as ~700 fM cDNA from HIV, 290 fM RNA from HCV, and 370 fM cDNA from SARS-CoV-2, all within 30 minutes without a need for target amplification. With isothermal amplification of SARS-CoV-2 RNA using RT-LAMP, the modified crRNAs were incorporated in a paper-based lateral flow assay that could detect the target with up to 23-fold higher sensitivity within 40-60 minutes.

## Main

Class 2 CRISPR/Cas (Clustered Regularly Interspaced Short Palindromic Repeats/CRISPR-associated proteins) systems, such as Cas12a (Cpf1, subtype V-A) and Cas13a (C2c2, subtype VI), are capable of nonspecific cleavage of ssDNA (single-stranded DNA) and RNA, respectively, in addition to successful gene editing.^1–3^ This attribute, known as trans-cleavage, is only activated once bound to an activator (ssDNA or dsDNA) that has complementary base-pairing to the guide crRNA. When combined with a FRET-based reporter, a fluorophore connected to a quencher via a short oligonucleotide sequence, the presence of the target activator can be confirmed. This phenomenon has been efficiently harnessed by SHERLOCK (Specific High-sensitivity Enzymatic Reporter unLOCKing) and DETECTR (DNA Endonuclease Targeted CRISPR Trans Reporter) to reliably detect nucleic acids.^1,4–8^

Research studies have reported that an extended secondary DNA on the guide crRNA for Cas12a or a hairpin RNA structure added to the sgRNA for Cas9 increases the efficiency and specificity of gene editing.^9,10^ In addition, chemically modified Cas12a guided-RNA has also been shown to facilitate improved gene correction in mammalian cells through both viral-and non-viral methods compared to the wild-type guide RNA.^11^ Though these modifications were employed to utilize the cis-cleavage aspect of the CRISPR/Cas systems, the effects of such alterations on the trans-cleavage remain unknown.

Based on the crystal structure of LbCas12a/crRNA/dsDNA (PDB ID: 5XUS)^12^, we reasoned that crRNA extensions can influence the trans-cleavage activity by either activating or inhibiting the catalytic efficiency of Cas12a, allowing us to better understand crRNA design with tunable trans-cleavage activity. We speculate that chemical modifications of the crRNA can potentially change its nature of binding and subsequently alter this collateral cleavage due to conformational changes of the Cas12a dynamic endonuclease domain.

We placed ssDNA, ssRNA, and phosphorothioate ssDNA extensions of various lengths ranging from 7 to 31 nucleotides on either the 3’- or 5’-ends of the crRNA targeting GFP (green fluorescent protein), referred to here as crGFP (Fig. 1b-h). In order to measure the collateral or trans-cleavage activity of Cas12a, we employed a FRET-based reporter used in DETECTR,^1^ composed of a fluorophore (FAM) and a quencher (3IABkFQ) connected by a 5-nucleotide sequence (TTATT), which displays increased fluorescence upon cleavage. Consistent with the previous literature^13^, when using wild-type crRNAs, we observed that the LbCas12a exhibited higher trans-cleavage activity than the AsCas12a or the FnCas12a, and therefore, we designed various modified crRNAs compatible with LbCas12a. Using the same reporters, we discovered that ssDNA and ssRNA extensions on the 3’-end of crGFP markedly enhanced the trans-cleavage ability of target-activated LbCas12a. Comparing the two types, the ssDNA extensions demonstrated higher activity than the corresponding ssRNA (Figs. 1b-d,f and Supplementary Figs. 1-5). On the other hand, the phosphorothioate ssDNA extensions at the 3’- or 5’-end displayed minimal or no activity, showing decreased fluorescence intensity as modification length increased (Figs. 1e,h and Supplementary Figs. 1-5). This observation suggests that further extending the crRNA with 13-mer phosphorothioate ssDNA and beyond significantly inhibits LbCas12a trans-cleavage activity. The finding corroborated B. Li and colleagues that phosphorothioate ssDNA may prevent crRNA-Cas12a-DNA complex formation.^14^

**Figure 1.**
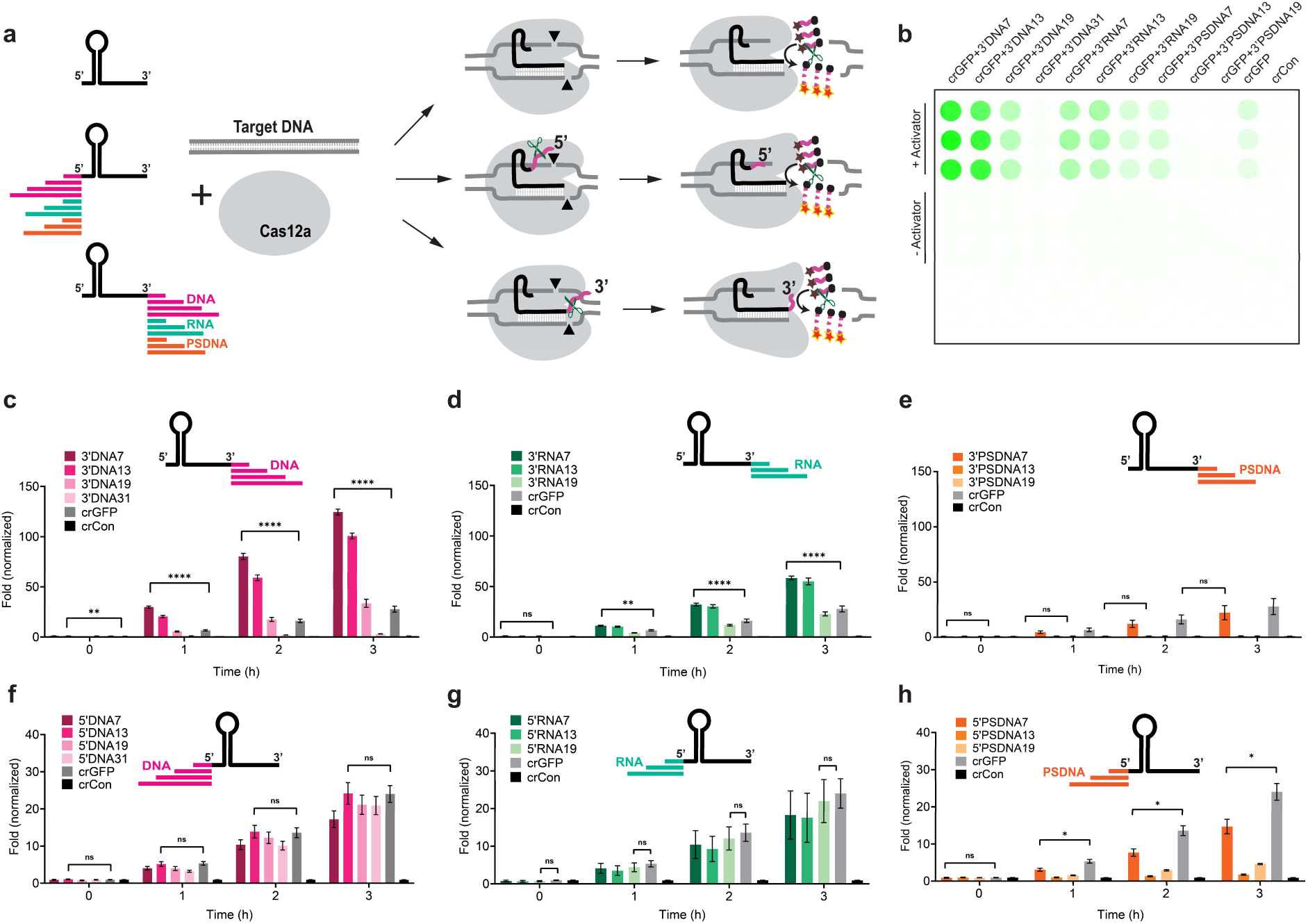
Trans-cleavage activity of LbCas12a with modified crRNA via fluorescence-quencher-based reporter assay with TA rich fluorophore-quencher systems tested. For other fluorophore-quencher systems, see supplementary figures 1-4. **(a)** Schematic diagram of cleavage of Cas12a with wild-type and modified crRNAs. The crRNA is extended on either the 3’- or 5’-ends with ssDNA, ssRNA, or phosphorothioate ssDNA. **(b)** A representation of a fluorescence-quencher-based trans-cleavage reporter assay image taken by GE Amersham Typhoon. **(c)**, **(d)**, and **(e)** 3’-end ssDNA, ssRNA and phosphorothioate ssDNA extensions of crRNA, respectively. **(f)**, **(g)**, and **(h)** 5’-end ssDNA, ssRNA and phosphorothioate ssDNA extensions of crRNA, respectively. The fold in fluorescence was normalized by taking the ratio of background-corrected fluorescence signals of sample with activator to the corresponding sample without activator. Error bars represent ± SEM, where n = 6 replicates; two-way ANOVA test two-way ANOVA (n=3, N=2, ^ns^P > 0.05, *P < 0.05, **P < 0.01, ***P < 0.001, ****P < 0.0001). The experiments were repeated at least twice with n = 3 per experiment.

Notably, the 3’-DNA with 7-mer extensions on the crGFP, referred as crGFP+3’DNA7, yielded the highest fluorescence signal compared to other modifications, measuring approximately 3.5-fold higher intensity than the wild-type crGFP (Fig. 1c, Supplementary Figs. 1a,2a). By investigating the conformational changes from the crystal structure of the binary LbCas12a:crRNA complex^12,15,16^, we observed that the 3’-end modifications on crRNA is proximal to the RuvC region of the LbCas12a. This supports our observation that the 3’-end extensions lead to higher trans-cleavage activity than the 5’-end. We speculated that once an R-loop is formed between crRNA and dsDNA or ssDNA activator, the LbCas12a executes a partial trans-cleavage of the 3’-end of crRNA, leaving an overhang. These remaining extensions may further expand the nuclease domain in the LbCas12a, resulting in conformational changes and allowing more access for nonspecific ssDNA cleavage. To confirm our hypothesis, we attached different fluorophores, or fluorophore-quencher pairs separated by DNA linkers, to either the 3’- or 5’-end of the crGFP with 7-mer DNA extensions and analyzed by denaturing gel electrophoresis. Surprisingly, we discovered that the 3’-end of the crRNA is processed by LbCas12a only in the presence of an activator while the 5’-end is cleaved by LbCas12a even in the absence of the activator (Fig. 2a,b and Supplementary Figs. 6,7).

**Figure 2:**
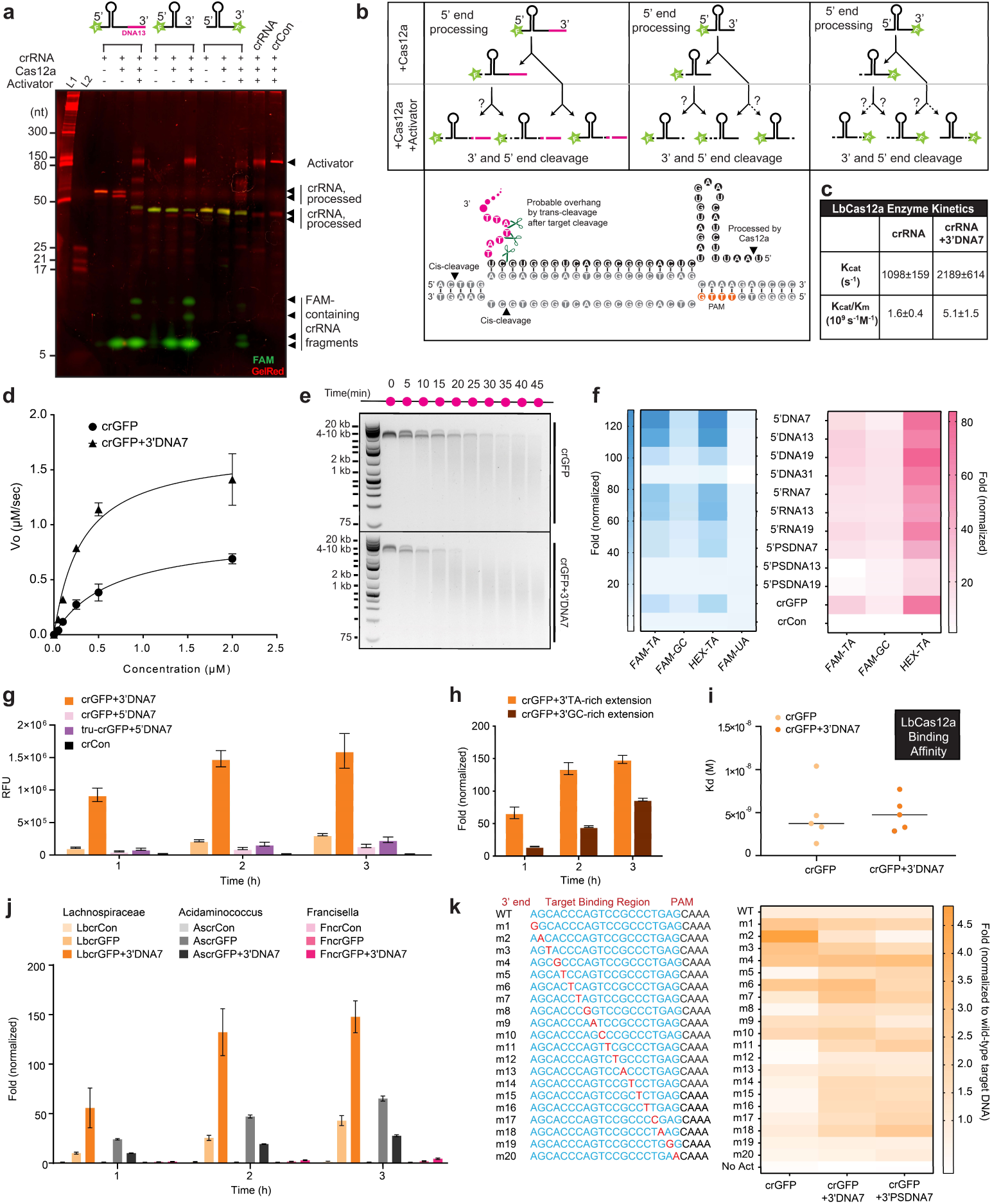
Mechanism and kinetics of LbCas12a trans-cleavage with modified crGFP. **(a)** Interactions of fluorescently labeled crRNAs with LbCas12a and dsDNA activator, characterized by PAGE analysis. In the absence of the activator, the modified crRNA (pre-crRNA) is trimmed by LbCas12a on its 5’-end (the first Uracil is cleaved, so-called truncated-crRNA or tru-crRNA). In the presence of the activator, the crRNA extensions are further trimmed, possibly leaving a 3’overhang. **(b)** Schematic diagram of putative processing of crRNA cleavage sites in the presence and absence of activator GFP. **(c)** schematic diagram of different cleavage sites in the presence and absence of activator GFP. **(c)** Enzyme kinetic data of LbCas12a with crGFP vs. crGFP+3’DNA7. **(d)** Michaelis-Menten kinetic study of the wild-type crGFP vs. crGFP+3’DNA7. For this assay, 100nM of LbCas12a, 100 nM of crRNA, and 7.4 nM of GFP fragment were used. **(e)** Time-dependent cis-cleavage of LbCas12a on GFP in the presence of nonspecific ssDNA M13mp18. The reaction mixture was taken out every five minutes and quenched with 6X SDS-containing loading dye. **(f)** Effect of different types of fluorophore-quencher systems on trans-cleavage activity with various modifications of crRNA. **(g)** Comparison of trans-cleavage activity between precursor crRNA (pre-crRNA) and mature crRNA (tru-crRNA, where the first Uracil on the 5’-end of the crRNA is cleaved by LbCas12a in the absence of the activator). **(h)** Comparison of trans-cleavage activity between AT-rich extensions and GC-rich extensions of the crRNA. **(i)** Dissociation constants of crGFP vs. crGFP+3’DNA7. The Kd was determined by the biolayer interferometry binding kinetic assay with R^2^>0.9. **(j)** Trans-cleavage activity of different variants of Cas12a. The prefix Lb, As, and Fn stand for Lachnospiraceae bacterium, Acidaminococcus, and Francisella novicida, respectively. **(k)** Single-point mutations (m1-m20) on the target strand of a dsDNA GFP activator. The heat map displays relative fluorescence intensity normalized to wild-type (WT) activator after 3 hours. Error bars represent ± SEM, where n = 6 replicates. The experiments were repeated at least twice with n = 3 per experiment.

By placing the fluorophore FAM on the 5’-end and a 7-mer DNA extension on the 3’-end of the crGFP, we learned that the first Uracil on the 5’-end of the crGFP gets trimmed by LbCas12a in the absence of an activator, which corroborated previous studies reported for FnCas12a^12^. As a result, the 5’-end modifications are eliminated and converted back to the wild-type crRNA before complexing with the activator. This finding reinforces our previous observation that the 5’-extended crRNA has similar collateral cleavage activity as the wild-type crRNA. Fascinated by LbCas12a pre-crRNA processing as previously described^17^ and from our observations, we investigated how extensions of the mature crRNA would influence the trans-cleavage activity compared to the corresponding extended pre-crRNA. We discovered that the modified pre-crRNA and modified mature crRNA (tru-crRNA) exhibited comparable trans-cleavage efficiency (Fig. 2g). Furthermore, when a dsDNA or an ssDNA activator was present, the 3’-and 5’-end DNA-extended crRNA were cleaved (Figs. 2a, 2b and Supplementary Figs. 6,7).

To further understand the LbCas12a enhanced enzymatic activity, we performed a Michaelis-Menten kinetic study on the wild-type crGFP and the crGFP+3’DNA7 and observed that the ratio Kcat/Km was 3.2-fold higher for crGFP+3’DNA7 than the unmodified crGFP (Figs. 2c,d). The time-dependent gel electrophoresis analysis of nonspecific cleavage of ssDNA M13mp18 phage (~7 kb) reconfirmed the fluorophore-quencher-based reporter assay results (Fig. 2e).

We speculated that the reporter composition itself may affect the LbCas12a collateral cleavage activity. Therefore, we incorporated and tested various nucleotides (GC and TA-rich) and fluorophores (FAM, HEX, and Cy5) within the reporter. Consistent with our hypothesis, we observed that the LbCas12a achieved maximal trans-cleavage activity with FAM or HEX and TA-rich reporter (Fig. 2f and Supplementary Figs. 1-5). Furthermore, these results led us to question whether the trans-cleavage activity is dependent on the sequence of ssDNA extensions on 3’-end of the crRNA. To test this, we altered the nucleotide content of the extended regions of the crGFP. It turned out that the crGFP with TA-rich extensions carried out significantly more collateral cleavage than those with GC-rich regions (Fig. 2h and Supplementary Fig. 8).

Based on our findings that the trans-cleavage activity is drastically improved by 7-mer ssDNA extensions to the 3’-end of crGFP, we questioned if the binding of crRNA with LbCas12a itself is influenced by such modifications. A biolayer interferometry binding kinetic assay revealed that the dissociation constant, K_d_, between the binary complex LbCas12a:crRNA and LbCas12a:crRNA+3’DNA7 are comparable within a low nM scale (Fig. 2i and Supplementary Fig. 9). These binding results suggest that the 3’DNA7 modification on crRNA does not affect the binary complex formation between the LbCas12a and the crRNA.

While 7-mer ssDNA extensions on the 3’-end of crRNA increases trans-cleavage activity with LbCas12a, we questioned if this is consistent across other orthologs of Cas12a. To investigate further, we carried out an in vitro cis-cleavage and trans-cleavage assay of AsCas12a and FnCas12a with an extended crGFP compared to a wild-type crGFP (Fig. 2j and Supplementary Fig. 10). Interestingly, the crGFP+3’DNA7 showed similar results with FnCas12a; however, it exhibited an opposite effect with AsCas12a. However, the cis-cleavage activity was found to be comparable between the crGFP and crGFP+3’DNA7 for all the orthologs tested. Overall, LbCas12a showed the highest fluorescence signal, which is consistent with previous studies.^13,18^ Through observation of the fluorophore-quencher-based reporter assay and time-dependent gel electrophoresis, we hypothesized that the various extensions of ssDNA on the crRNA induce conformational changes on LbCas12a that result in enhanced endonuclease activity.

Structural analysis of LbCas12a shows that it contains a single RuvC domain, which processes precursor crRNA into mature crRNA, cleaves target dsDNA or ssDNA (referred here as activators), and executes nonspecific cleavage afterwards.^19,20^ Therefore, we were interested in understanding the effects of these modified crRNAs on cis-cleavage compared to the wild-type crRNA, as well as how cis-cleavage activity correlates to the trans-cleavage activity. Towards this, we carried out an in vitro cis-cleavage assay for various 3’-end and 5’-end modifications. We noticed that the cis-cleavage activity was either similar or marginally improved with most 3’-end modifications while the 5’-end modifications showed either similar or slightly reduced activity. This phenomenon suggests that the trans-cleavage activity is commensurate with the cis-cleavage activity (Supplementary Figs. 11-12).

Next, we sought to the characterize specificity of these extended crRNAs in discriminating point mutations across dsDNA. By mutating a single nucleotide at each position across the target-binding region, we discovered that the crGFP+3’DNA7 tolerated mutations and produced a stronger fluorescence signal than the wild-type crGFP (Supplementary Fig.13). However, the fluorescence intensity ratio of mutated to the non-mutated dsDNA targets for the crGFP+3’DNA7 was quite comparable to the crGFP (Fig. 2k and Supplementary Fig.13). This observation suggests that the modified crRNAs increased sensitivity of trans-cleavage, however, the specificity remained unchanged.

Previous studies demonstrated that FnCas12a is a metal-dependent endonuclease, and magnesium ions are required for FnCas12a-mediated self-processing of precursor crRNA.^20^ Based on these findings, we hypothesized that different metal ions may significantly affect the trans-cleavage activity of LbCas12a. This led us to test a range of divalent metal cations and discovered that most ions including Ca^2+^, Co^2+^, Zn^2+^, Cu^2+^, and Mn^2+^ significantly inhibited the LbCas12a activity (Supplementary Fig. 14). By further investigating the Zn^2+^ mediated inhibition of LbCas12a, we found that the inhibition was dose-dependent (Supplementary Fig. 15). Interestingly, Ni^2+^ ions showed an unusual cis-cleavage activity possibly due to its interactions with the His tags on LbCas12a (Supplementary Fig. 14). Among the tested divalent metal ions, the Mg^2+^ ions showed the highest in vitro cis-cleavage activity, which was consistent with the literature.^20^ Therefore, we characterized the effect of Mg^2+^ ions on trans-cleavage activity of LbCas12a. With increasing the concentration of Mg^2+^ ions, a significant increase in fluorescence signal was observed in an in vitro trans-cleavage assay. By varying the amount of Mg^2+^ in the Cas12a reaction, we identified that the optimal condition of Mg^2+^ was around 13 mM (Fig. 3a-b and Supplementary Figs. 16-19).

**Figure 3.**
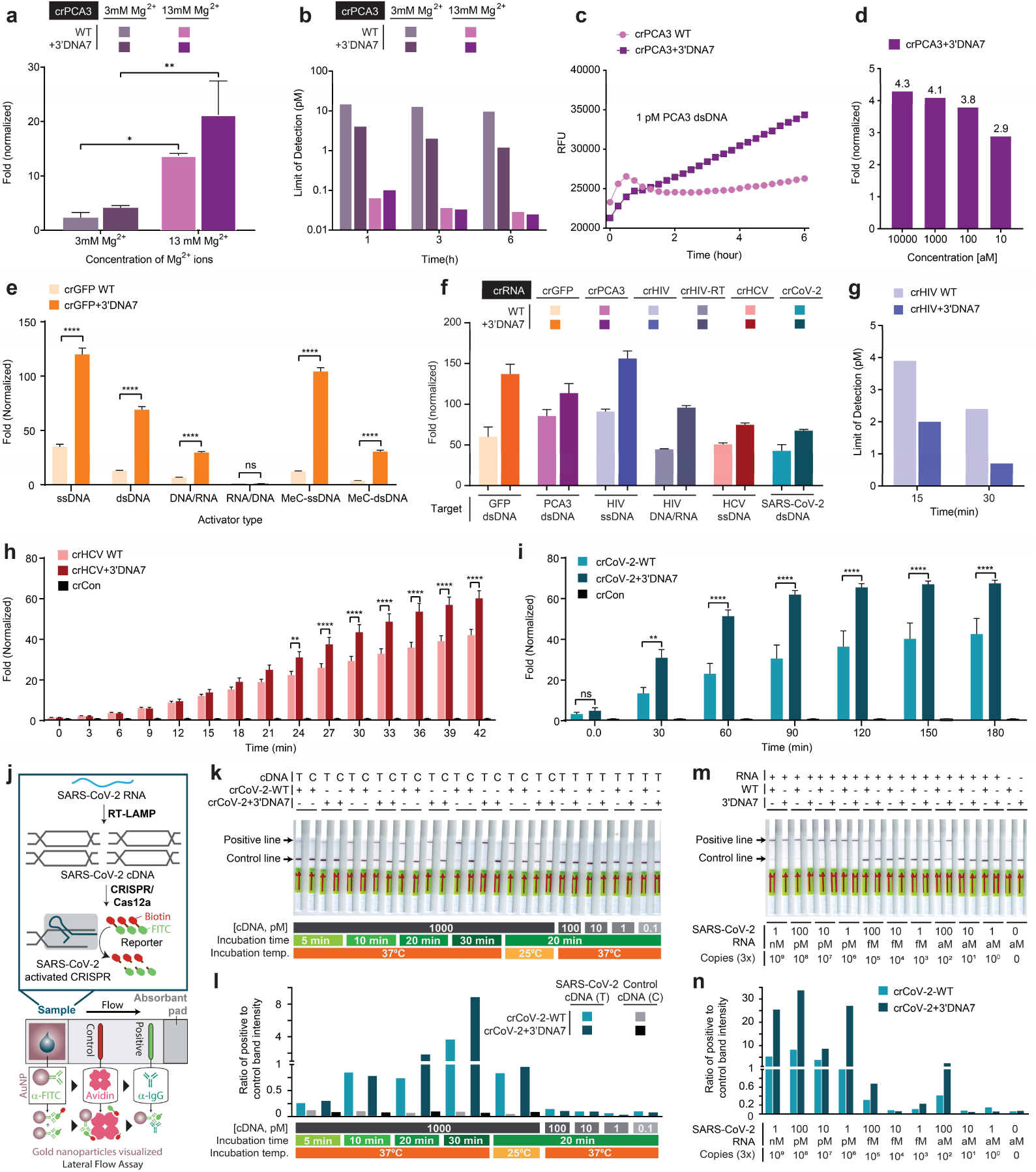
Improved detection of various targets using modified crRNA/LbCas12a system. **(a)** Effect of Magnesium ion on the trans-cleavage activity. **(b)** Limit of detection in femtomolar targeting PCA3 in simulated human urine at optimized Mg^2+^ concentration. **(c)** Raw fluorescence data showing detectable signal difference at 1 pM of PCA3 DNA. **(d)** Fold change in fluorescence signal of the modified crRNA+3’DNA7 targeting GFP after the recombinase polymerase amplification (RPA) step. **(e)** Effect of heteroduplex DNA-RNA and methylated activators on the trans-cleavage activity of LbCas12a. **(f)** trans-cleavage activity of different DNA targets. GFP, PCA3, COVID-19, and HCV are dsDNA activators, and their fluorescence shown were taken after 3 hours. The HIV target is ssDNA or cDNA/RNA heteroduplex, and its fluorescence signal shown was taken after 1 hour. **(g)** Limit of detection targeting HIV cDNA fragment after 15 and 30 minutes. Results in **(b)** and **(g)** are based on limit of detection calculations. **(h)** Fold change in trans-cleavage activity with LbCas12a in presence of 100 pM (10 fmols) of target HCV ssDNA. Using modified crRNA, the limit of detection of HCV target ssDNA was found to be 290 fM (29 amoles) at 30 min, without target amplification, mean ± SE, two-way ANOVA (n=3, N=3). **(i)** Fold change in trans-cleavage activity with LbCas12a in presence of 100 pM (10 fmols) of target SARS-CoV-2 cDNA (dsDNA), mean ± SE, two-way ANOVA (n=3, N=2, ^ns^P > 0.05, *P < 0.05, **P < 0.01, ***P < 0.001, ****P < 0.0001). **(j)** Lateral flow assay detecting 1 nM (100 fmols) of SARS-CoV-2 cDNA using either crCoV-2 or crCoV-2+3’DNA-7 within 20 minutes of incubation. **(j)** Schematic diagram showing how a lateral flow assay works. Briefly, the dipstick uses gold-labeled FITC-specific antibodies that binds to FITC-biotin reporter and travel through membrane. Only cleaved reporter will reside at the positive line. **(k)** Lateral flow assay detecting SARS-CoV-2 cDNA using crCov-2 and crCoV-2+3’DNA7 without a pre-amplification step, and **(l)** band-intensity analysis of (k) using ImageJ. **(m)** Lateral flow assay detecting SARS-CoV-2 RNA N gene using crCov-2 and crCoV-2+3’DNA7 with RT-LAMP, and **(n)** band-intensity analysis of (m) using ImageJ.

We optimized the previously developed CRISPR-based detection assays^1,5,6^ and combined them with our engineered crRNA+3’DNA7 to create a CRISPR-ENHANCE (Enhanced analysis of nucleic acids with crRNA extensions) technology or referred here as ENHANCE. To validate the ENHANCE technology, we first selected a clinically relevant nucleic acid biomarker, Prostate Cancer Antigen 3 (PCA3/DD3), which is one of the most overexpressed genes in prostate cancer tissue and excreted in patients’ urine. Consequently, elevated PCA3 levels during prostate cancer progression has become a widely targeted biomarker for detection.^21–24^ To determine the limit of detection of PCA3 using our ENHANCE technology, we spiked the PCA3 cDNA into synthetic urine and investigated how this clinically-relevant environment affects the activity of Cas12a.

Using ENHANCE for detecting the PCA3 cDNA, the limit of detection was determined to be as low as 25 fM in the urine at 13 mM Mg^2+^ concentration compared to ~1 pM at 3 mM Mg^2+^ concentration after 6 hours (Figs. 3a-c and Supplementary Figs. 17-19). In contrast, the wild-type crRNA also showed a similar 29 fM limit of detection at 13 mM Mg^2+^ concentration while the limit of detection was ~10 pM at 3 mM Mg^2+^ concentrations after 6 hours. Therefore, by combining the crRNA modifications with increased Mg^2+^ ion concentrations, we achieved approximately 400-fold increase in sensitivity, based on limit of detection calculations. Nevertheless, this observation also suggests that our modified crRNA+3’DNA7 significantly improves the limit of detection at low Mg^2+^ but reaches a saturation point that is comparable with the wild-type crRNA at high Mg^2+^ concentration. To understand the importance of divalent ions in the Cas12a trans-cleavage reaction, we carried out a Michaelis-Menten kinetic study with various Mg^2+^ concentrations (Supplementary Fig. 17). We observed that the initial reaction rate of Cas12a in the presence of high Mg^2+^ concentrations increased tremendously compared to that in low Mg^2+^.

However, the two reaction rates eventually reach a similar saturation point (Supplementary Figs. 16-18). This suggests that Mg^2+^ is not only required for the Cas12a reaction, but also accelerates the enzyme’s trans-cleavage activity. Regardless, Mg^2+^ plays an important role in lowering the limit of detection in synthetic urine containing PCA3. While as low as 25 fM (equivalent to 2.5 attomoles) of PCA3 cDNA can be detected with ENHANCE without any target amplification (Supplementary Fig. 19), the clinical concentration of PCA3 mRNA in the urine can be lower and therefore may require target pre-amplification.^25,26^ Therefore, we incorporated and tested a recombinase polymerase amplification (RPA) step to isothermally amplify the PCA3 cDNA. By combining the RPA step as previously reported,^1,7^ the concentration of PCA3 cDNA in the urine was detectable down to ~10 aM (1 zmol) with 2.9-fold signal to noise ratio (Fig. 3d).

While crRNA/LbCas12a has been traditionally used to detect unmodified DNA, the field is missing the knowledge on how the common epigenetic marker, DNA methylation, affects its trans-cleavage activity. DNA methylation is also one of the bacterial defense systems that fight against outside invaders. It would be fascinating to understand how LbCas12a collateral cleavage is able to recognize methylated DNA targets. This curiosity let us to discover that the wild-type crRNA had significantly reduced activity in detecting methylated DNA, containing 5-methyl cytosine, compared to the unmethylated DNA. However, the ENHANCE showed 5.4-fold and 3.4-fold and higher trans-cleavage activity compared to the wild-type crRNA for targeting the methylated dsDNA and ssDNA, respectively (Fig. 3e, Supplementary Fig. 20a).

Although there are no reports on RNA-guided RNA targeting by LbCas12a, we envisioned that an RNA can potentially be detected as a DNA/RNA heteroduplex. To test this hypothesis, we incorporated a reverse transcription step to convert RNA into cDNA/RNA heteroduplex before detecting the RNA with a trans-cleavage assay. We discovered that the RNA can only be detected if the target strand for crRNA is a DNA but not an RNA in a heteroduplex. Notably, the efficiency of the trans-cleavage activity for the DNA/RNA heteroduplex was found to be significantly lower than the corresponding ssDNA or dsDNA (Fig. 3e, Supplementary Fig. 20b). However, the DNA/RNA heteroduplex achieved an improved enzymatic collateral activity when using the crRNA+3’DNA7 compared to the wild-type crRNA. We applied the ENHANCE to successfully detect low picomolar concentrations of HIV RNA target encoding Tat gene with our DNA/RNA heteroduplex detection strategy (Fig. 3f). In parallel, ssDNA and dsDNA targets from HIV were also detected with much higher sensitivity compared to the wild-type crRNA within 15 to 30 minutes (Figs. 3f,g and Supplementary Figs. 21). We further applied the ENHANCE for detecting HCV ssDNA and HCV dsDNA gene encoding a polyprotein precursor, both of which indicated consistent enhanced collateral activity than the wild-type crRNAs within 24 minutes (Figs. 3f, 3h, and Supplementary Fig. 22). The limit of detection for HIV and HCV targets were calculated to be 700 fM cDNA and 290 fM ssDNA, respectively.

In the wake of the recent COVID-19 pandemic, there is an urgent need to rapidly detect the SARS-CoV-2 coronavirus (referred as CoV-2 here for simplicity). We optimized the ENHANCE to detect CoV-2 dsDNA by designing crRNAs targeting nucleocapsid phosphoprotein encoding N gene (Figs. 3f,i). While no clinical samples were tested, the results indicated the 3’DNA7-modified crRNA consistently demonstrated higher sensitivity for detecting CoV-2 dsDNA within 30 minutes as compared to the wild-type crCoV-2 (Supplementary Figs. 23-25). By incorporating a commercially available paper-based lateral flow assay with a FITC-ssDNA-Biotin reporter,^27–29^ we could visually detect 1 nM of CoV-2 cDNA within 20 minutes of incubation using both wild-type and modified crRNAs without any target amplification (Fig. 3j-l and Supplementary Fig. 26). The enzyme trans-cleavage activity exhibited a consistent trend with the crRNA+3’DNA7 among five different targets (Fig. 3f). When incorporating a reverse transcription step and a loop-mediated isothermal amplification (RT-LAMP) strategy into the ENHANCE, both the crCoV-2-WT and the crCoV-2+3’DNA7 demonstrated a limit of detection down to a 3-300 copies of RNA (Fig. 3m). However, in case of crCoV-2-WT, the partial cleavage of the reporter resulted in a darker control line on the paper strip. Band-intensity analysis showed that the ENHANCE exhibited an average of 23-fold higher ratio of positive to control line between 1 nM and 1 pM of target CoV-2 RNA, while the crCoV-2-WT indicated an average of only 7-fold ratio (Figs. 3m,n and Supplementary Fig. 27). Additionally, the time lapse pictures of lateral flow assays showed that the positive line for target-containing samples developed and became visible sooner (within 30 seconds) for the crCoV-2+3’DNA7 than the crCoV-2-WT (Supplementary Fig. 28).

In summary, we extended the 3’-and 5’-end of the crRNA and discovered an amplified trans-cleavage activity of LbCas12a when the 3’-end is extended with DNA or RNA. We applied this modified crRNA/LbCas12a system with the optimal conditions to detect PCA3 in simulated urine with high sensitivity. This ENHANCE technology enabled us to detect the DNA/RNA heteroduplex and methylated DNA with unprecedented sensitivity. We further employed this system to test a range of target nucleic acids, including ssDNA, dsDNA, and RNA from HIV, HCV, and SARS-CoV-2 without the need for further optimization. These findings are a crucial step towards enhancing detection of nucleic acids and assisting in the diagnosis of various diseases.

## Methods

### DNA activator preparation

Multiple DNA activators were used in this study. The GFP fragment (942 bp) was produced by amplifying the pEGFP-C1 plasmid using polymerase chain reaction in the Proflex PCR system (ThermoFisher Scientific). The PCR product was purified using Monarch® Nucleic Acid Purification Kit (New England Biolabs Inc.).

Additionally, the 40-nt ds-GFP and ds-PCA3 activators were generated by annealing two single-stranded TS and NTS fragments at a 1:1 ratio (Integrated DNA Technologies Inc.) in 1X hybridization buffer (20 nM Tris-Cl, pH 7.8, 100 mM KCl, 5 mM MgCl_2_). The annealing process was executed in the Proflex PCR system at 90oC for 2 minutes followed by gradual cooling to 25oC at a rate of 0.1 °C/s.

### LbCas12a expression and purification

The plasmid LbCpf1-2NLS (Addgene #102566, a gift from Jennifer Doudna Lab)^30^ was transformed into Nico21(DE3) competent cells (New England Biolabs). Colonies were picked and inoculated in Terrific Broth at 37°C until OD600 = 0.6. IPTG was then added to the cultures, and they were grown at 18°C overnight.

Cell pellets were collected from the overnight cultures by centrifugation, resuspended in lysis buffer (2M NaCl, 20 mM Tris-HCl, 20 mM imidazole, 0.5 mM TCEP, 0.25 mg/ml lysozyme, and 1mM PMSF, PH = 8), and broken by sonication. The sonicated solution then underwent high speed centrifugation at 40,000 RCF for 45 minutes. The collected supernatant was then run through a Ni-NTA Hispur column (Thermofisher) pre-equilibrated with wash buffer A (2M NaCl, 20 mM Tris-HCl, 20 mM imidazole, 0.5 mM TCEP, PH = 8). The column was then eluted with buffer B (2M NaCl, 20 mM Tris-HCl, 300 mM imidazole, 0.5 mM TCEP, PH = 8). The eluted fractions were then pooled together and underwent TEV cleavage overnight at 4°C (TEV protease was purified using the plasmid pRK793, #8827 from Addgene, a gift from David Waugh Lab).

The resulting fraction was equilibrated with buffer C (100 mM NaCl, 20 mM HEPES, 0.5 mM TCEP, PH = 8) at a 1:1 ratio and run through Hitrap Heparin HP 1 ml column (GE Biosciences). The column was washed with buffer C and gradually eluted at a gradient rate with buffer D (100 mM NaCl, 20 mM HEPES, 0.5 mM TCEP, PH = 8). The eluted fraction was concentrated down to 500 μL and passed through the Hiload Superdex 200 pg column (GE Biosciences). The purified LbCas12a was then buffer exchanged with storage buffer (500 mM NaCl, 20 mM Na_2_CO_3_, 0.1mM TCEP, 50% glycerol, PH = 6) and flash frozen at −80°C until use.

### Biolayer Interferometry (BLI) binding kinetic assay

The BLI Ni-NTA biosensors were purchased from Fortebio to perform the binding kinetic study with polyhistidine-tagged LbCas12a. In detail, the experiment was carried out in a 96-well plate and included five steps: baseline, loading, baseline2, association, and dissociation. The biosensors were dipped into the baseline containing 1X kinetic buffer (1X PBS, 0.1% BSA, and 0.01% Tween 20). They were then transferred into each loading well containing 10 μg/ml of LbCas12a. After processing through loading and baseline2, the protein-tagged biosensor was next allowed to dip into the crRNA sample wells at different dilution (10, 5, 2.5, 1.25, 0.625, 0.3125, 0.15625, and 0 μg/ml) in the association step. The dissociation step occurred when the biosensors were transferred back to baseline2 at a shake speed of 1000 rpm. All the samples were read by the Octet QKe system (Fortebio). Kd was determined by software Data Analysis 10.0 (Fortebio), and only Kd with R^2^>0.9 were extracted for comparison between crRNA wild type and modified crRNAs.

### Cis-cleavage assay

In-vitro digestion reactions were carried out with three different types of the Cas12a family (LbCas12a, AsCas12a, and FnCas12a were purchased from New England Biolabs Inc., Integrated DNA Technologies Inc., and abm®, respectively) and a wide array of modified crRNA’s (purchased from DNA Technologies Inc.). Cas12a and crRNA were mixed with 1:1 ratio (100 nM:100 nM) in 1X NEBuffer 2.1 and pre-incubated at 25°C for 15 minutes to promote the ribonucleoprotein complex formation. DNA activator (GFP or PCA3 fragments) (final concentration of 7 nM) was then added to the mixture to produce a total volume of 30 μL and incubated at 37°C for 30 minutes. The sample was then analyzed in either 1% agarose gel (for GFP fragments amplified from the pEGFP-C1 plasmid) pre-stained with either SYBER Gold (Invitrogen), GelRed (Biotium Inc.), or premade 15% Novex™ TBE-Urea Gel (Invitrogen)^31^.

### M13mp18 nonspecific cleavage assay

Nonspecific cleavage activity of Cpf1 was activated by incubating Cpf1, crRNA, and DNA activator with a concentration of 100 nM:100 nM: 2nM in 1X NEBuffer 2.1 buffer at 37 °C for 30 minutes. M13mp18 was then added to the 30 μL reaction mixture and incubated for an additional 45 minutes. A fraction of the reaction was taken out every 5 minutes, quenched in 6X purple gel loading dye (New England Biolabs Inc.), and subsequently analyzed in 1% agarose gel (Fisher Scientific)^1^.

### Trans-cleavage reporter assay

The fluorophore-quencher reporter assay was carried out following a standard clinical detection protocol. The Cas12a-crRNA ribonucleoprotein complex was assembled by mixing 100 nM Cas12a and 100 nM crRNA in 1X NEBuffer 2.1 in the Proflex PCR system (ThermoFisher Scientific) at 25°C for 15 minutes (volume 28.5 µL). The activator (1 nM final concentration), FQ reporter (500 nM final concentration), and UltraPure™ DNase/RNase-Free distilled water (Invitrogen) were pre-added to a 96-well plate (Greiner Bio-One) to a volume of 71.5 µL. The reaction was initiated by adding the Cas12a-crRNA mixture to the 96-well plate preloaded with activator and FQ reporter (Integrated DNA Technologies Inc). The plate was quickly transferred to a plate reader (ClarioStar or BioTek), and fluorescence intensity was measured every 3 minutes for 3 or 12 hours (detection limit assay) (FAM FQ: λ_ex_: 483/30 nm, λ_em_: 530/30 nm; HEX: λ_ex_: 430/20 nm, λ_em_: 555/30 nm). After 3 or 12 hours (detection limit assay), the sample was scanned for images using the Amersham Typhoon (GE Healthcare).

For Michaelis-Menten kinetic study, 30 nm LbCas12a: 30 nM crRNA: 1 nM activator were mixed in NEBuffer 2.1 and incubated at 37°C for 30 minutes. The reaction mixture was then transferred to a 96-well plate (Greiner Bio-One) preloaded with different concentrations of FQ reporter (HEX or FAM FQ reporter: 0 M, 0.05 μM, 0.1 μM, 0. 25 μM, 0.5 μM, and 1 μM) and UltraPure™ DNase/RNase-Free distilled water (Invitrogen)^1^.

To find limit of detection (LoD), the fluorophore-quencher reporter assay was carried out with various concentrations of activator. The LoD calculations were based on the following formula^32^:

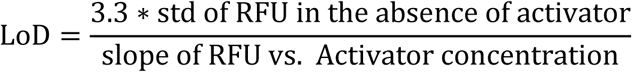

### Effects of metal ions on Cas12a cleavage study

The metal ions (Mg^2+^, Zn^2+^, Mn^2+^, Cu^2+^, Co^2+^, Ca^2+^) were prepared by diluting chloride salt in different concentrations. For cis-cleavage, the Cas12a-crRNA-metal iron duplex was mixed with 100 nM: 100nM: varying nM ratio in 1X annealing buffer (100 mM Tris-HCl, pH 7.9 @ 25°C, 500 mM NaCl, 1 mg/ml BSA) and pre-incubated at 25°C for 15 minutes. DNA activator (GFP or PCA3 fragments) was then added to the mixture to a total volume of 30 μL and incubated at 37°C for 30 minutes.

### Paper Strip Test

To minimize the testing time, the following reagent was assembled in a one-pot reaction:

a. 10X NEBuffer: 3 µL (0.3X final concentration)
b. 3 µM LbCas12a: 1 µL (300 nM final concentration)
c. 3 µM crRNA: 2 µL (600 nM final concentration)
d. 5 µM FAM-biotin reporter: 3 µL (500 nM final concentration)
e. Various concentration of Activator: 2 µL
f. Nuclease-free water: 89 µL (total reaction volume = 100 µL)

The reaction mixture was incubated at either 37°C or 25°C for different time periods (5, 10, and 20 minutes). A Milenia HybriDetect (TwistDx) strip was dipped in each reaction and allowed for rapid visualization.

For experiment involving a recombinase polymerase amplification (RPA) step, the reaction mix was prepared in the following order:

a. Forward primer (10 µM): 2.4 µL
b. Reverse primer (10 µM): 2.4 µL
c. Primer free rehydration buffer: 29.5 µL
d. Template and nuclease-free water: 13.2 µL
e. (CH_3_COO)_2_Mg (280 mM): 2.5 µL (total volume = 50 µL)

The RPA reaction was incubated at 39°C for 20-30 minutes prior to LbCas12a reaction.

For experiments involving a RT-LAMP preamplification step of target RNA, the mixture was prepared in the following order (except for the RNA and primer mix samples (IDT Technologies), everything was purchased from New England Biolabs):

a. 10X isothermal amplification buffer: 2.5 µL
b. 100 mM MgSO_4_: 1 µL
c. 10 mM dNTP: 3.5 µL
d. 10X primer mix (2 µM F3, 2 µM B3, 16 µM FIP, 16 µM BIP, LF and BF are optional): 2.5 µL
e. Bst 2.0 polymerase: 1 µL
f. Warmstart RT master mix: 1 µL
g. RNase inhibitor, murine (NEB # M0314S): 1 µL
h. Nuclease-free water: 7.5 µL
i. RNA sample: 5 µL (total volume = 25 µL)

The RT-LAMP reaction was incubated at 65°C for 20-30 minutes prior to LbCas12a reaction.

## Competing interests

The authors declare no competing interests.

## Author contributions

PKJ initiated the study; LTN and PKJ designed research; LTN and BMS performed research; LNT, BMS, and PKJ analyzed the data; LN and PKJ wrote the manuscript that was edited and approved by all authors.

## Acknowledgements

We are grateful to the members in the Jain lab for their helpful discussions and the University of Florida (UF) Health Cancer Center for their support. We are particularly thankful to Eric Beck for editing the manuscript and Ling Jin, Santosh Rananaware, and Marco Downing for helping with the experiments and/or data analysis. We also thank the Monoclonal Antibody core facility staff, especially Dr. Angle Sampson and Shadi Bootorabi, at the UF Interdisciplinary Center for Biotechnology Research (ICBR) for coordinating the biolayer interferometry experiments. This research was supported by the internal funding from the UF and the UF Herbert Wertheim College of Engineering.

## Notes

### Competing Interest Statement

The authors have declared no competing interest.

## REFERENCES

1. Chen, J. S. et al. CRISPR-Cas12a target binding unleashes indiscriminate single-stranded DNase activity. Science (2018). doi:10.1126/science.aar6245

2. Wang, Q. et al. The CRISPR-Cas13a Gene-Editing System Induces Collateral Cleavage of RNA in Glioma Cells. Adv. Sci. 6, (2019).

3. Li, S. Y. et al. CRISPR-Cas12a has both cis- and trans-cleavage activities on single-stranded DNA. Cell Res. 28, 491–493 (2018).

4. Gootenberg, J. S. et al. Nucleic acid detection with CRISPR-Cas13a/C2c2. Science 356, 438–442 (2017).

5. Gootenberg, J. S. et al. Multiplexed and portable nucleic acid detection platform with Cas13, Cas12a and Csm6. Science (2018). doi:10.1126/science.aaq0179

6. Harrington, L. B. et al. Programmed DNA destruction by miniature CRISPR-Cas14 enzymes. Science 362, 839–842 (2018).

7. Kellner, M. J., Koob, J. G., Gootenberg, J. S., Abudayyeh, O. O. & Zhang, F. SHERLOCK: nucleic acid detection with CRISPR nucleases. Nat. Protoc. 14, 2986–3012 (2019).

8. Abudayyeh, O. O., Gootenberg, J. S., Kellner, M. J. & Zhang, F. Nucleic Acid Detection of Plant Genes Using CRISPR-Cas13. Cris. J. 2, 165–171 (2019).

9. Kocak, D. D. et al. Increasing the specificity of CRISPR systems with engineered RNA secondary structures. Nat. Biotechnol. (2019). doi:10.1038/s41587-019-0095-1

10. Park, H. M. et al. Extension of the crRNA enhances Cpf1 gene editing in vitro and in vivo. Nat. Commun. 9, 1–12 (2018).

11. McMahon, M. A., Prakash, T. P., Cleveland, D. W., Bennett, C. F. & Rahdar, M. Chemically Modified Cpf1-CRISPR RNAs Mediate Efficient Genome Editing in Mammalian Cells. Mol. Ther. 26, 1228–1240 (2018).

12. Yamano, T. et al. Structural Basis for the Canonical and Non-canonical PAM Recognition by CRISPR-Cpf1. Mol. Cell 67, 633–645.e3 (2017).

13. Fuchs, R. T., Curcuru, J., Mabuchi, M., Yourik, P. & Robb, G. B. Cas12a trans-cleavage can be modulated in vitro and is active on ssDNA, dsDNA, and RNA. bioRxiv 600890 (2019). doi:10.1101/600890

14. Li, B. et al. Synthetic Oligonucleotides Inhibit CRISPR-Cpf1-Mediated Genome Editing. Cell Rep. 25, 3262–3272.e3 (2018).

15. Yamano, T. et al. Crystal Structure of Cpf1 in Complex with Guide RNA and Target DNA. Cell 165, 949–962 (2016).

16. Dong, D. et al. The crystal structure of Cpf1 in complex with CRISPR RNA. Nature 1–16 (2016). doi:10.1038/nature17944

17. Swarts, D. C., van der Oost, J. & Jinek, M. Structural Basis for Guide RNA Processing and Seed-Dependent DNA Targeting by CRISPR-Cas12a. Mol. Cell 66, 221–233.e4 (2017).

18. Dai, Y. et al. Exploring the Trans-Cleavage Activity of CRISPR-Cas12a (cpf1) for the Development of a Universal Electrochemical Biosensor. Angew. Chemie - Int. Ed. 58, 17399–17405 (2019).

19. Zetsche, B. et al. Cpf1 Is a Single RNA-Guided Endonuclease of a Class 2 CRISPR-Cas System. Cell 163, 759–771 (2015).

20. Fonfara, I., Richter, H., BratoviÄ, M., Le Rhun, A. & Charpentier, E. The CRISPR-associated DNA-cleaving enzyme Cpf1 also processes precursor CRISPR RNA. Nature 532, 517–521 (2016).

21. Marks, L. S. & Bostwick, D. G. Prostate Cancer Specificity of PCA3 Gene Testing: Examples from Clinical Practice. Rev. Urol. 10, 175–81 (2008).

22. Laxman, B. et al. A first-generation multiplex biomarker analysis of urine for the early detection of prostate cancer. Cancer Res. (2008). doi:10.1158/0008-5472.CAN-07-3224

23. Yang, Z., Yu, L. & Wang, Z. PCA3 and TMPRSS2-ERG gene fusions as diagnostic biomarkers for prostate cancer. Chinese J. Cancer Res. 28, 65–71 (2016).

24. Filella, X. et al. PCA3 in the detection and management of early prostate cancer. Tumor Biology (2013). doi:10.1007/s13277-013-0739-6

25. Ferro, M. et al. Prostate Health Index (Phi) and Prostate Cancer Antigen 3 (PCA3) Significantly Improve Prostate Cancer Detection at Initial Biopsy in a Total PSA Range of 2-10 ng/ml. PLoS One 8, 1–7 (2013).

26. Loeb, S. & Partin, A. W. PCA3 Urinary Biomarker for Prostate Cancer. Rev. Urol. 12, e205–6 (2010).

27. Gootenberg, J. S. et al. Multiplexed and portable nucleic acid detection platform with Cas13, Cas12a and Csm6. Science 360, 439–444 (2018).

28. Zhang, F., Abudayyeh, O. O., Gootenberg, J. S., Sciences, C. & Mathers, L. A protocol for detection of COVID-19 using CRISPR diagnostics v.20200321. Sherlock Biosciences, Broad Institute, MIT: Cambridge, MA (2020).

29. Broughton, J. P. et al. A protocol for rapid detection of the 2019 novel coronavirus SARS-CoV-2 using CRISPR diagnostics: SARS-CoV-2 DETECTR v3. Mammoth Biosciences, South San Francisco, CA (2020).

30. Moreno-Mateos, M. A. et al. CRISPR-Cpf1 mediates efficient homology-directed repair and temperature-controlled genome editing. Nat. Commun. 8, 1–9 (2017).

31. Zetsche, B. et al. Cpf1 Is a Single RNA-Guided Endonuclease of a Class 2 CRISPR-Cas System. Cell 163, 759–771 (2015).

32. Hajian, R. et al. Detection of unamplified target genes via CRISPR–Cas9 immobilized on a graphene field-effect transistor. Nat. Biomed. Eng. 3, 427–437 (2019).

