## Supplementary Information for "Enhancement of trans-cleavage activity of Cas12a with engineered crRNA enables amplified nucleic acid detection"

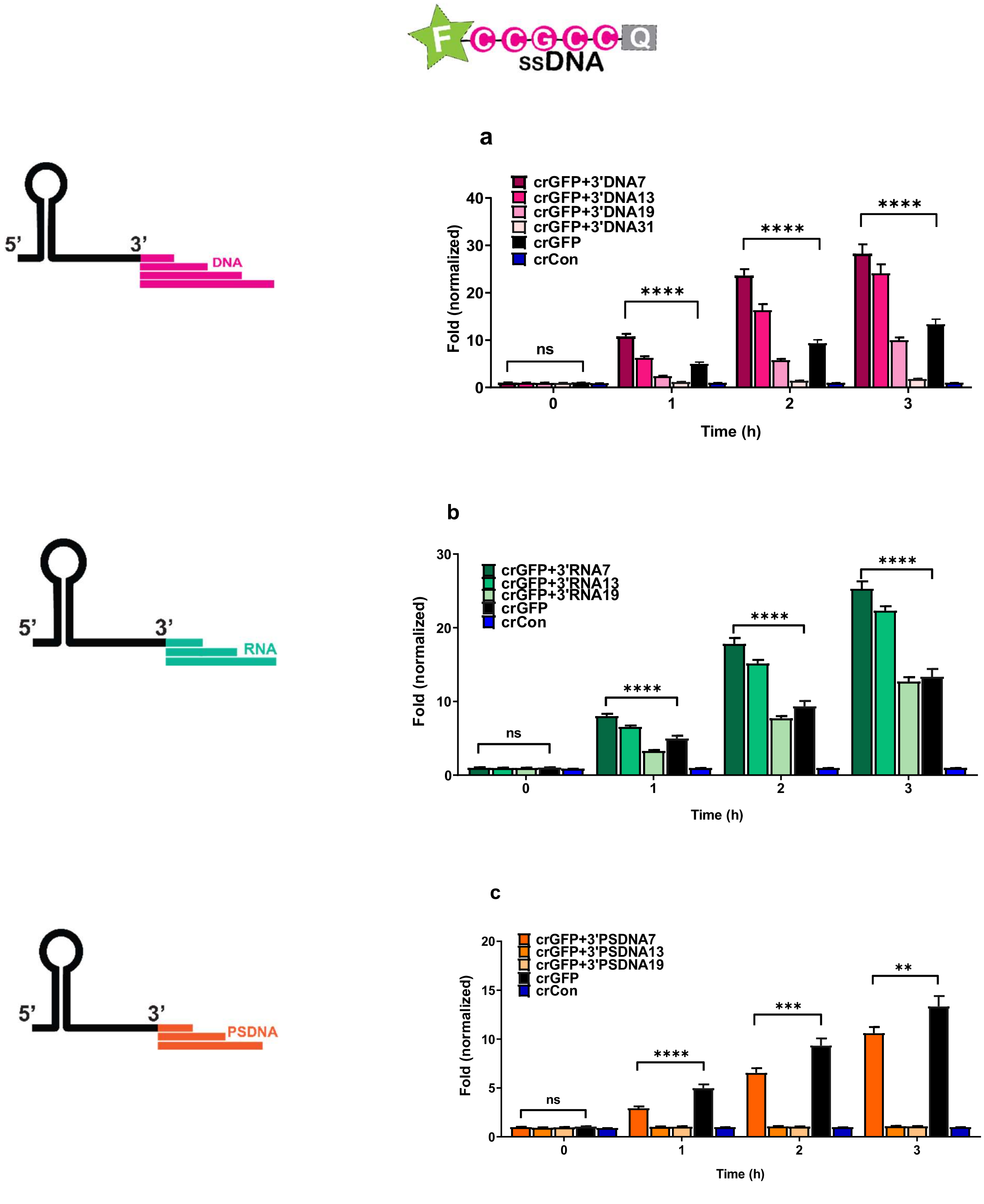

**Supplementary Figure 1**. Trans-cleavage activity of LbCas12a with modified 3' crRNA via fluorescence-quencher reporter assay with FAM-GC, where FAM-GC is a reporter shown above**.** F stands for fluorophore, and Q stands for quencher. (**a**) DNA, (**b**) RNA, and (**c**) PSDNA extensions of crRNA. The fold in fluorescence was normalized by taking the ratio of background-corrected fluorescence signals of the samples with the activator to the corresponding samples without the activator. Error bars represent ± SEM, where n = 6 replicates; two-way ANOVA test (n=3, N=2, $^{ns}P > 0.05$, $*P < 0.05$, $**P < 0.01$, $***P < 0.001$, $****P < 0.0001$). The experiments were repeated at least twice with n = 3 per experiment.

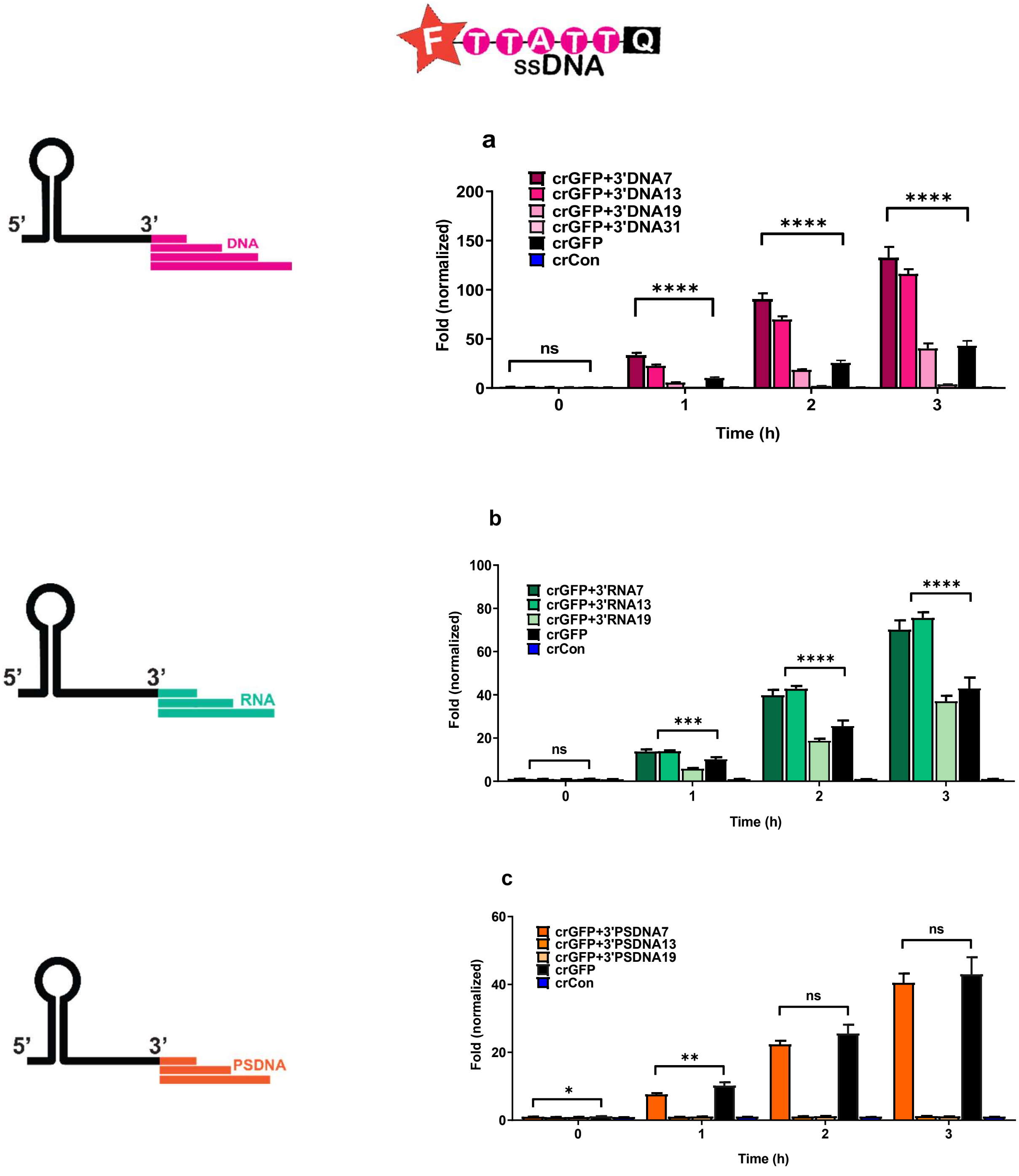

**Supplementary Figure 2**. Trans-cleavage activity of LbCas12a with modified 3' crRNA via fluorescence-quencher reporter assay with HEX-TA, where HEX-TA is a reporter shown above. F stands for fluorophore, and Q stands for quencher. (**a**) DNA, (**b**) RNA, and (**c**) PSDNA extensions of crRNA. The fold in fluorescence was normalized by taking the ratio of background-corrected fluorescence signals of the samples with the activator to the corresponding samples without the activator. Error bars represent ± SEM, where n = 6 replicates; two-way ANOVA test (n=3, N=2, $^{ns}P > 0.05$, $*P < 0.05$, $**P < 0.01$, $***P < 0.001$, $****P < 0.0001$). The experiments were repeated at least twice with n = 3 per experiment.

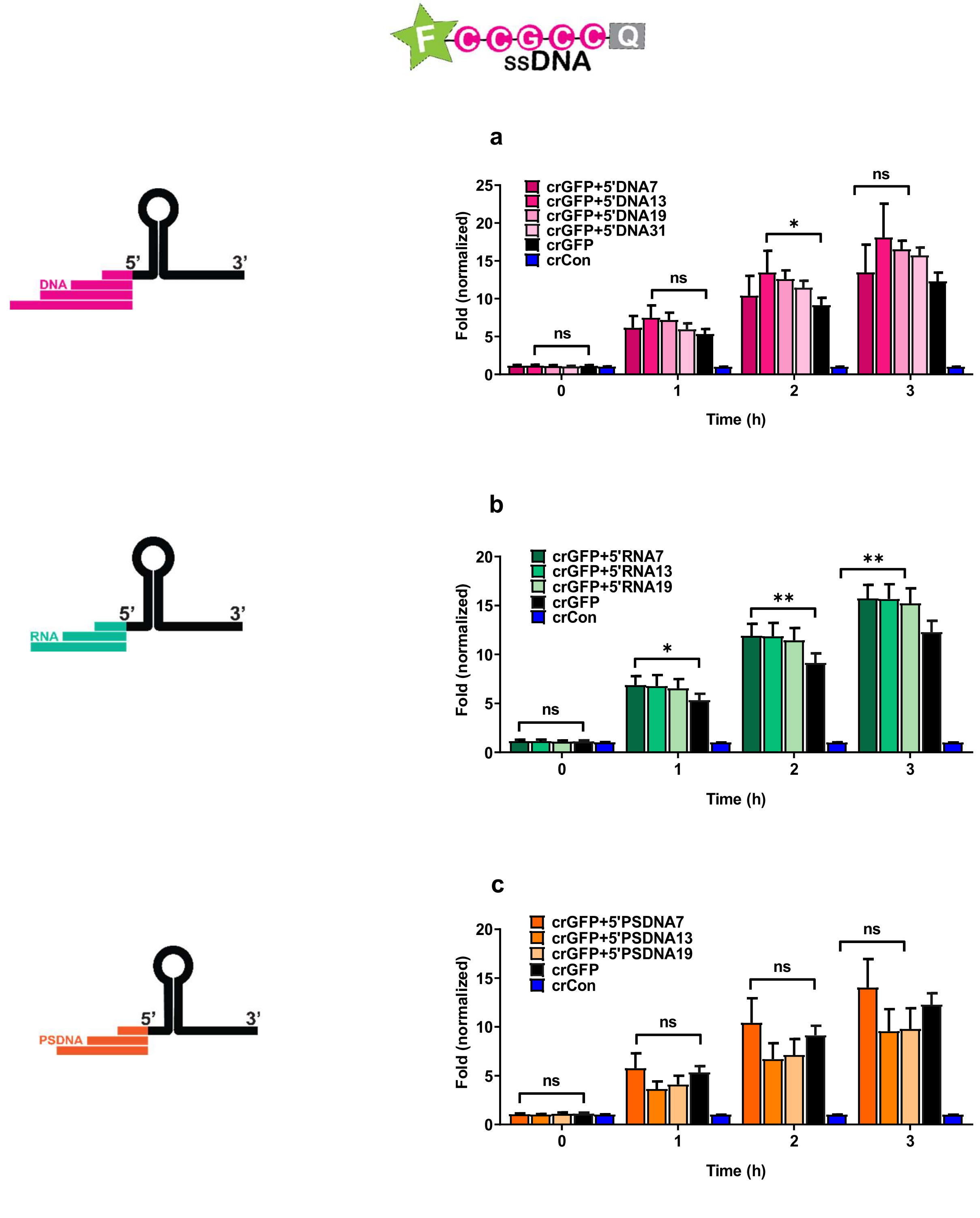

**Supplementary Figure 3.** Trans-cleavage activity of LbCas12a with modified 5’ crRNA via fluorescence-quencher reporter assay with FAM-GC, where FAM-GC is a reporter shown above. F stands for fluorophore, and Q stands for quencher. (**a**) DNA, (**b**) RNA, and (**c**) PSDNA extensions of crRNA. The fold in fluorescence was normalized by taking the ratio of background-corrected fluorescence signals of the samples with the activator to the corresponding samples without the activator. Error bars represent ± SEM, where n = 6 replicates; two-way ANOVA test (n=3, N=2, $^{ns}P > 0.05$, $^{*}P < 0.05$, $^{**}P < 0.01$, $^{***}P < 0.001$, $^{****}P < 0.0001$). The experiments were repeated at least twice with n = 3 per experiment.

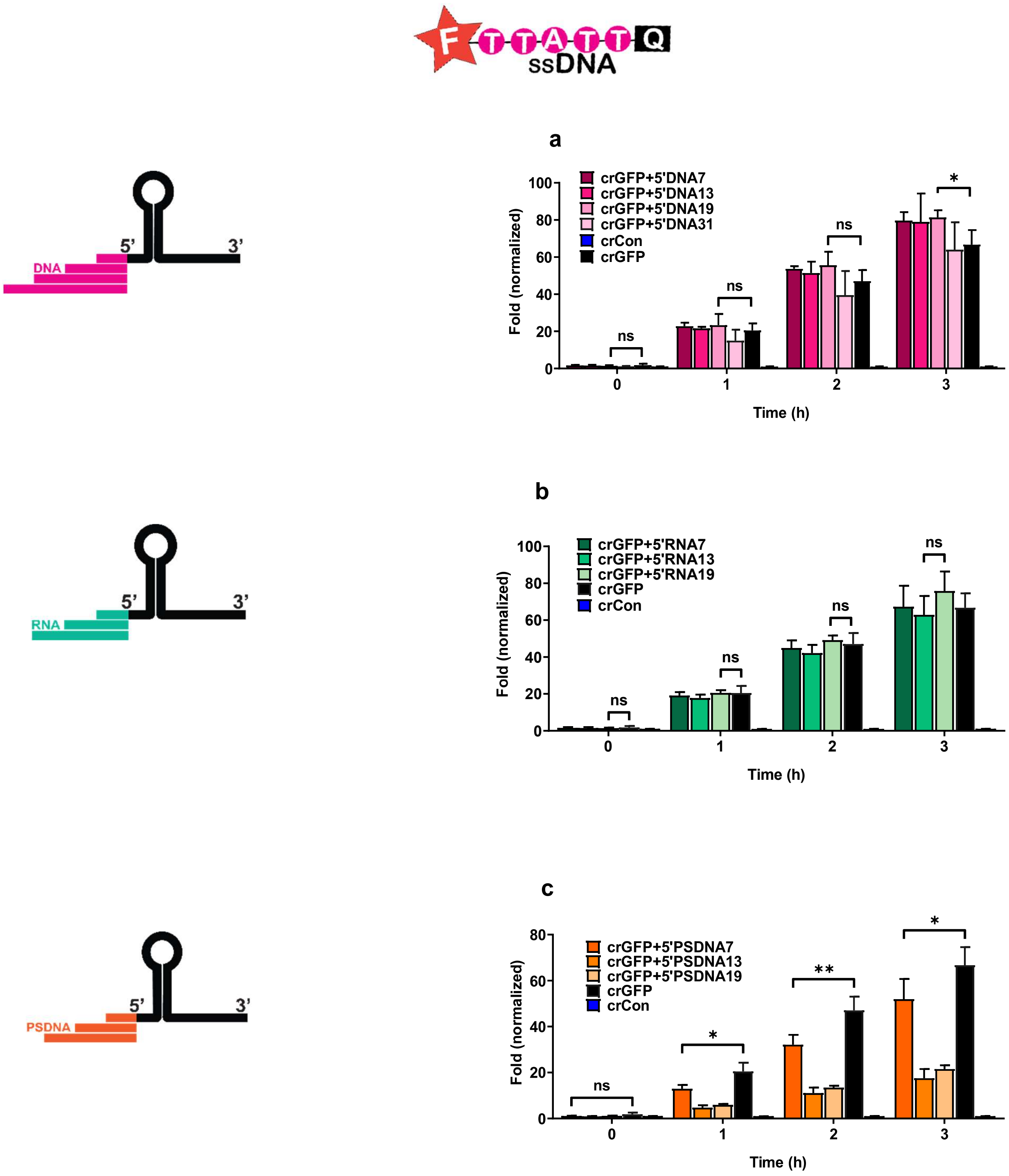

**Supplementary Figure 4.** Trans-cleavage activity of LbCas12a with modified 5' crRNA via fluorescence-quencher reporter assay with HEX-TA, where HEX-TA is a reporter shown above. F stands for fluorophore, and Q stands for quencher. (**a**) DNA, (**b**) RNA, and (**c**) PSDNA extensions of crRNA. The fold in fluorescence was normalized by taking the ratio of background-corrected fluorescence signals of the samples with the activator to the corresponding samples without the activator. Error bars represent ± SEM, where n = 6 replicates; two-way ANOVA test (n=3, N=2, $^{ns}P > 0.05$, $^{*}P < 0.05$, $^{**}P < 0.01$, $^{***}P < 0.001$, $^{****}P < 0.0001$). The experiments were repeated at least twice with n = 3 per experiment.

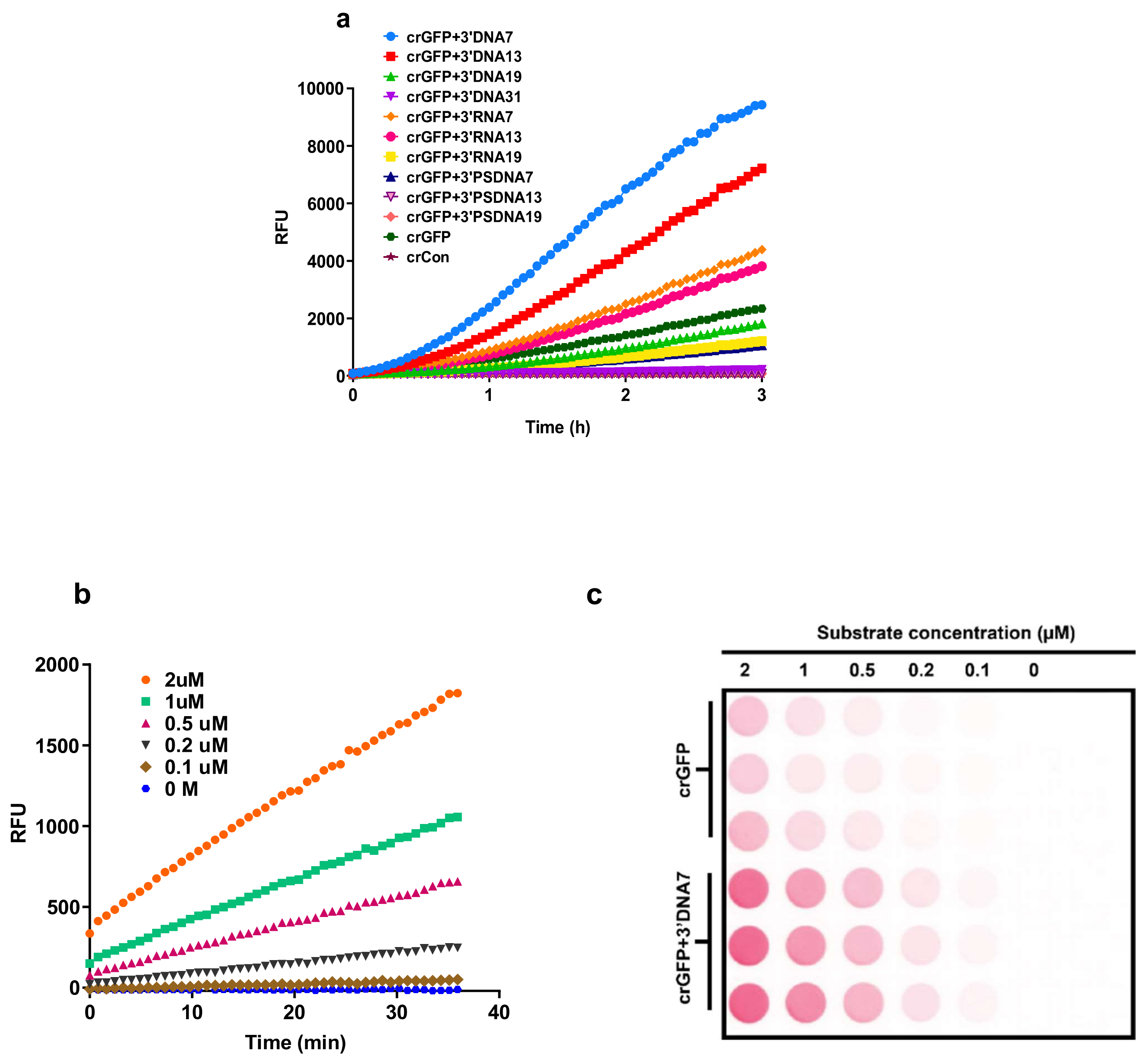

**Supplementary Figure 5.** Representatives of samples from the fluorescence-quencher reporter assay. (**a**) Fluorescence measurement in relative fluorescence unit (RFU) of LbCas12a reaction with 3'-end modified crGFP. (**b**) Fluorescence reading of the Michaelis-Menten study. The substrate used in this experiment was /5HEX/TTA TT/3IABkFQ/. (**c**) Fluorescence image of (b) using the Amersham Typhoon.

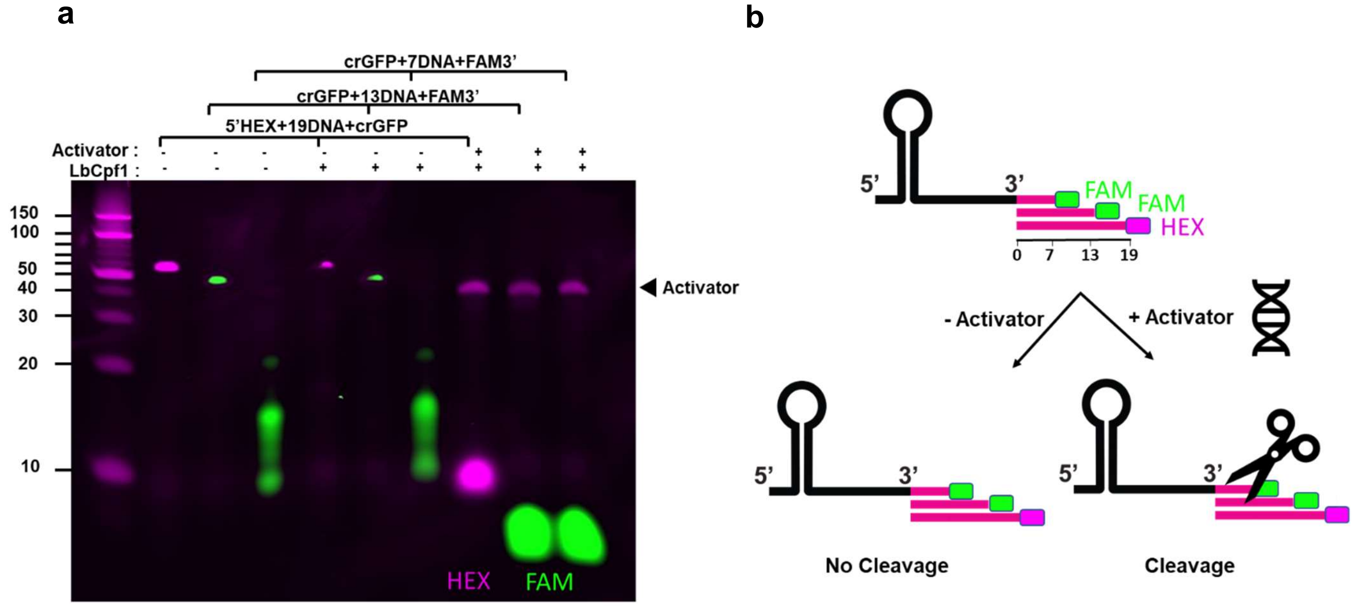

**Supplementary Figure 6.** Cis-cleavage of LbCas12a with fluorescently-labeled extended crGFP. (**a**) The 3'- end extensions of crGFP were varied from 0 to 19-mer DNA and tagged with FAM/HEX. (**b**) The proposed mechanism for LbCas12a processing of modified crGFP as observed from (a).

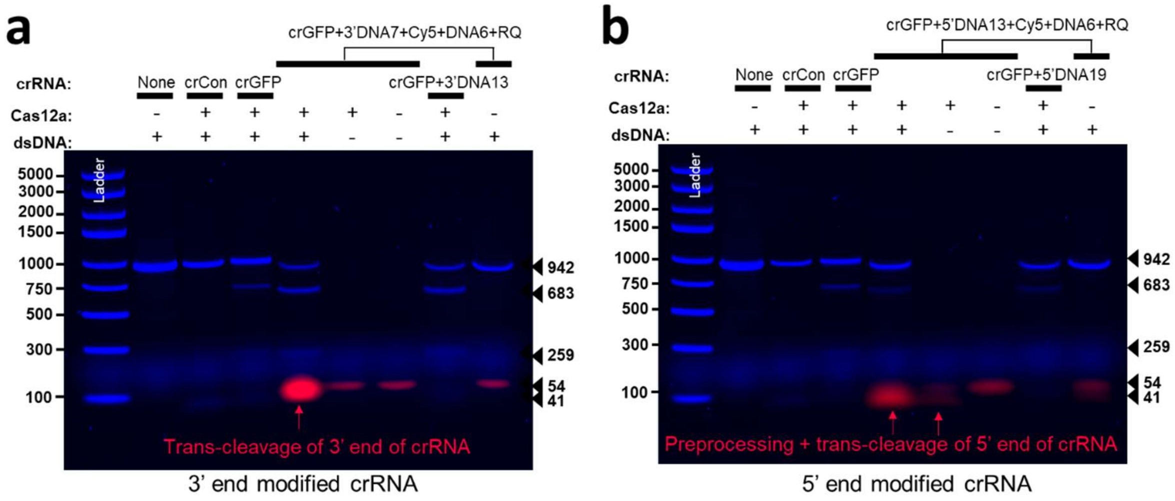

**Supplementary Figure 7**. Gel images of cis-cleavage assay of LbCas12a with different crRNAs carrying a fluorophore-quencher pair on either (a) 3' end or (b) 5' end. crCon (scrambled crRNA), crGFP (GFP targeting crRNA), (a) crGFP+3' DNA13, and crGFP+3' DNA7+Cy5+DNA6+Iowa Black RQ or (b) 5'-end modified crRNAs including crGFP+5' DNA19 and crGFP+5' DNA13+Cy5+DNA6+Iowa Black RQ. Cy5 is indicated in red and DNA stained with GelRed is shown in blue. The 5' end modified crRNAs showed cleavage of crRNA immediately after adding the LbCas12a but before adding the activator. However, both 3' and 5' end modified crRNAs, showed increase in signal intensity after activator addition indicating trans cleavage of the crRNA. 250 nM of LbCpf1, 250 nM of crRNA, and 7.4 nM of DNA activator fragment were used.

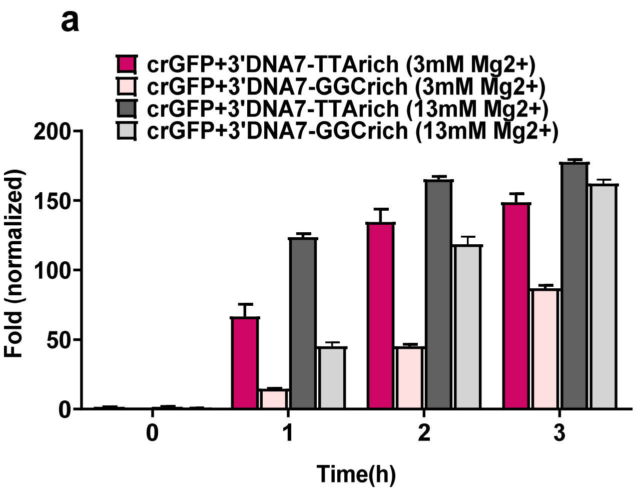

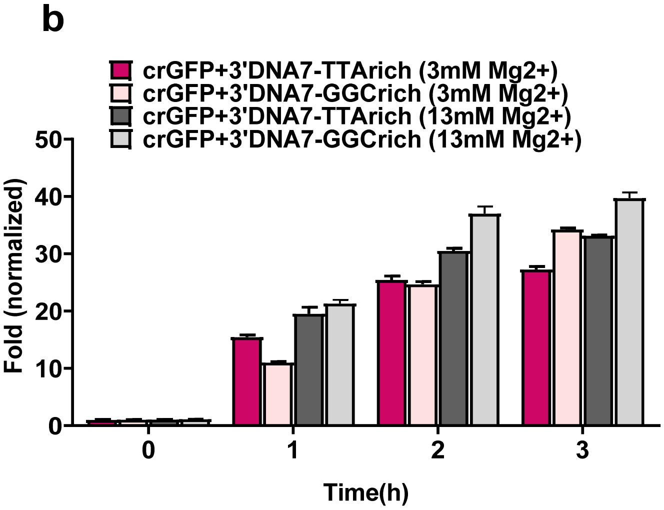

**Supplementary Figure 8.** Trans-cleavage activity of LbCas12a with modified crGFP+3'DNA7 with either GC rich or TA rich region via fluorescence-quencher reporter assay at varying $Mg^{2+}$ concentration**.** F stands for fluorophore, and Q stands for quencher. (**a**) FAM-TA was used, (**b**) FAM-GC was used. The fold in fluorescence was normalized by taking the ratio of background-corrected fluorescence signals of the samples with the activator to the corresponding samples without the activator. Error bars represent ± SEM, where n = 6 replicates. The experiments were repeated at least twice with n = 3 per experiment.

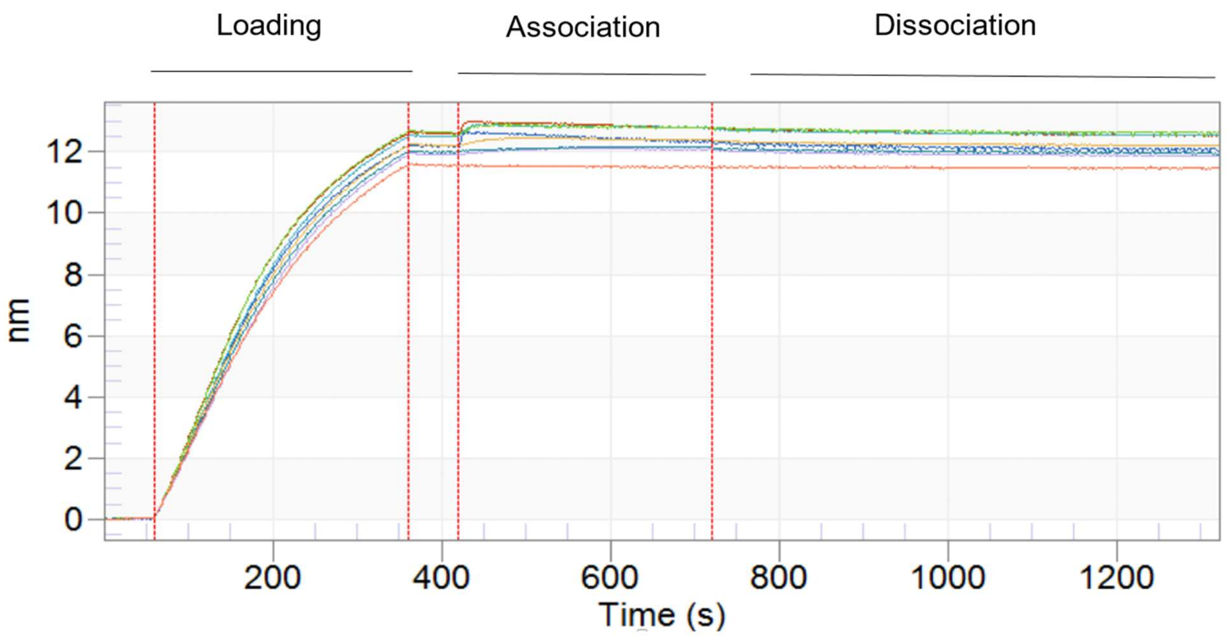

**Supplementary Figure 9.** A representation of the biolayer interferometry (BLI) binding kinetics. The picture shown above is the binding kinetic study of LbCas12a to crGFP+3'DNA7. The experiment was carried in five steps: baseline1, loading, baseline2, association, and dissociation (see materials and methods). The y-axis represents response of LbCas12a and crRNA to the biosensor in nm. Data were trimmed and processed using the Octet Data Analysis 10.0 software, and only $K_D$ values with $R^2 > 0.9$ were selected.

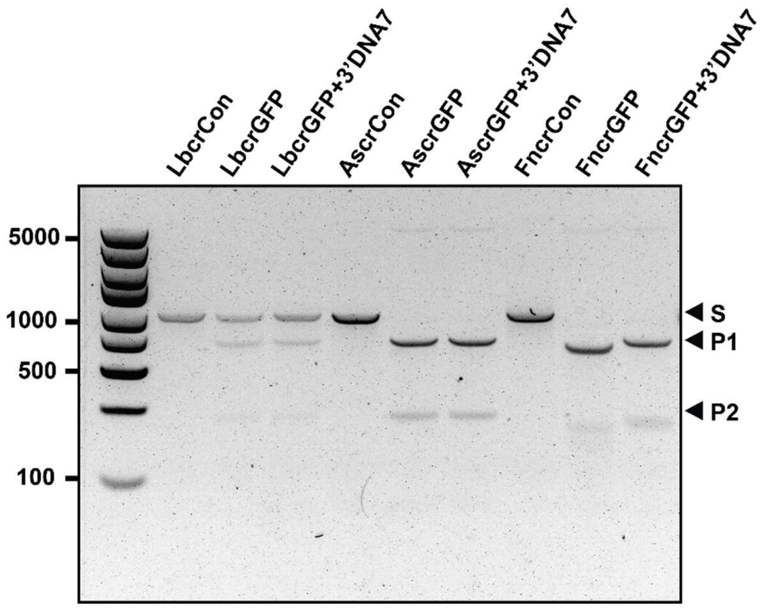

**Supplementary Figure 10.** Cis-cleavage and trans-cleavage activity of LbCas12 with modified crGFP and crGFP+3'DNA7 with LbCas12a, AsCas12a, and FnCas12a**.** The cis-cleavage reaction was loaded on 1% native agarose gel.

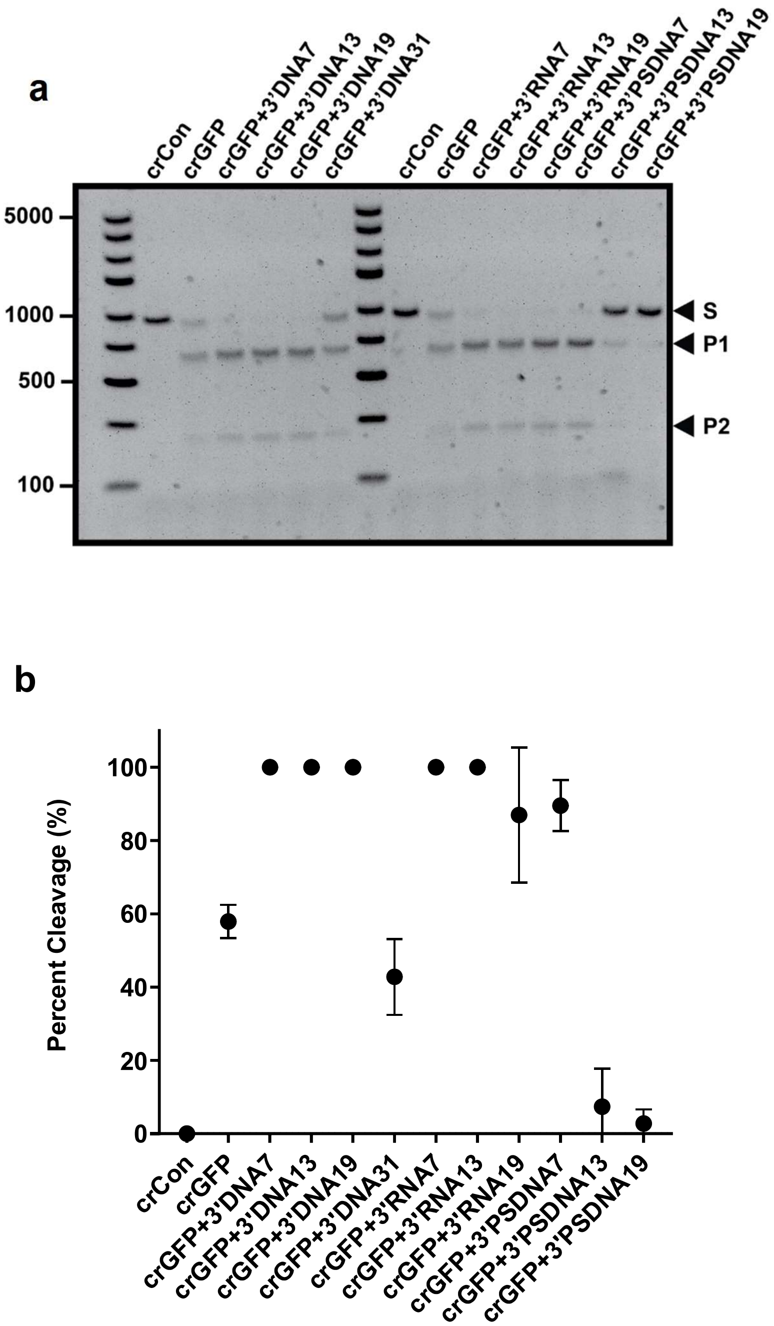

**Supplementary Figure 11.** Cis-cleavage of LbCas12a with 3'-end modified crGFP. For this assay, 100nM of LbCas12a, 100 nM of crRNA, and 7.4 nM of GFP fragment were used. crCon represents nonspecific crRNA. (**a**) 1% agarose gel image. (**b**) Percent cleavage of the GFP fragment calculated in (**a**). The experiment was repeated more than twice.

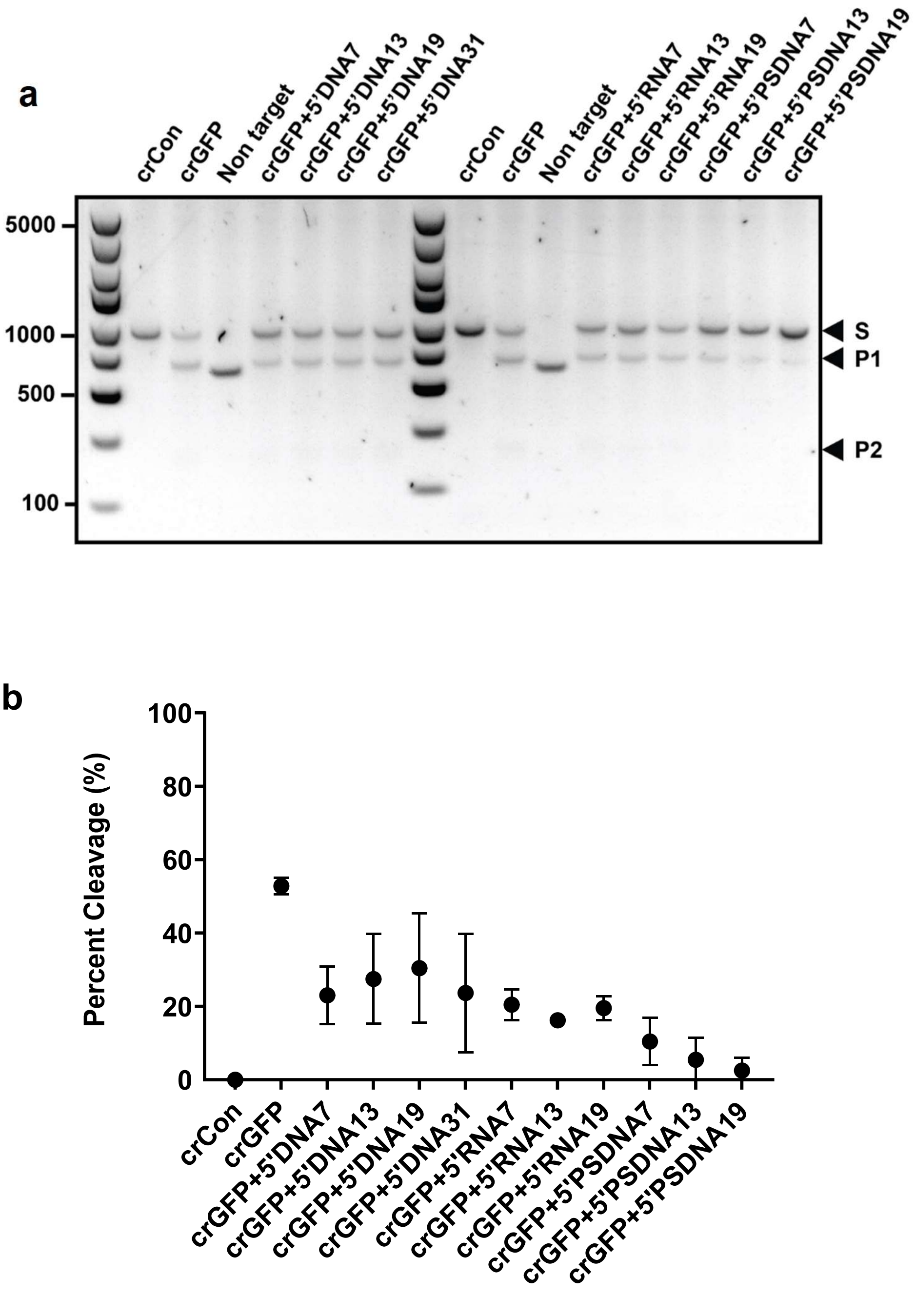

**Supplementary Figure 12.** Cis-cleavage of LbCas12a with 5'-end modified crGFP. For this assay, 100nM of LbCas12a, 100 nM of crRNA, and 7.4 nM of GFP fragment were used. crCon represents nonspecific crRNA. (**a**) 1% agarose gel image. (**b**) Percent cleavage of the GFP fragment calculated in (**a**). The experiment was repeated more than twice.

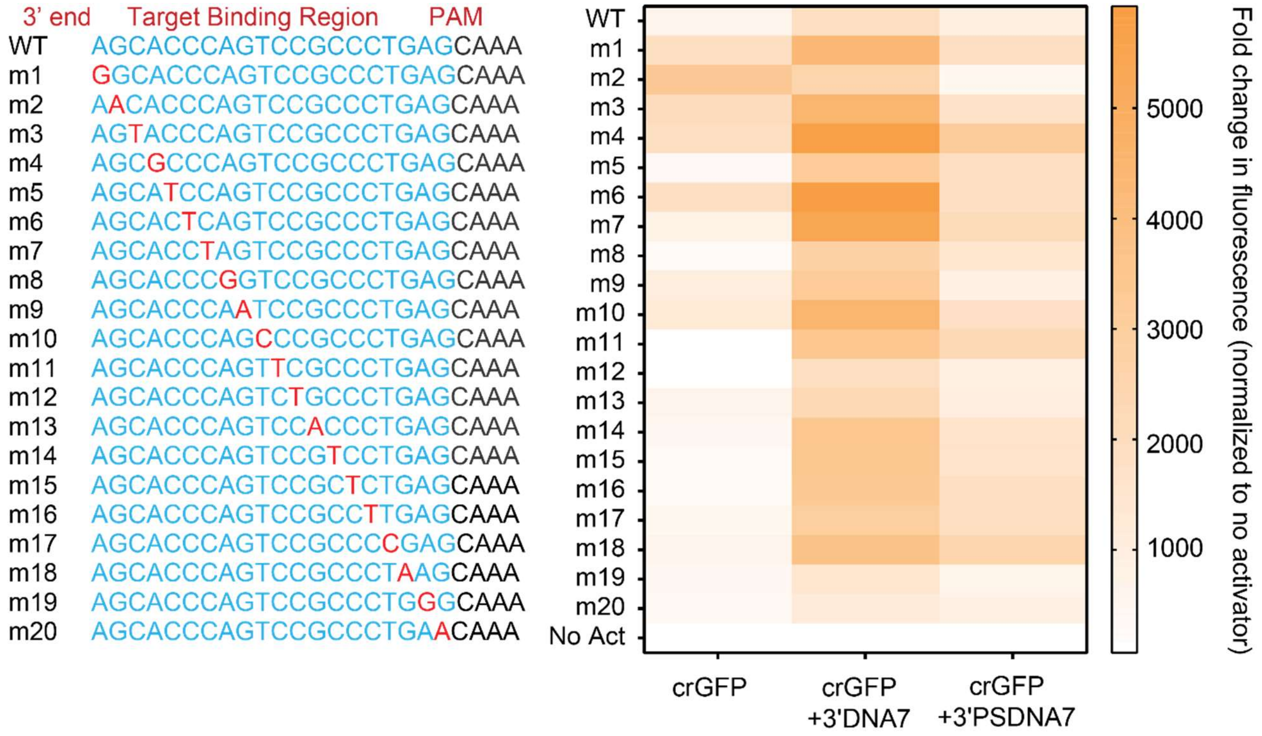

**Supplemental Figure 13.** Single-point mutation on the target strand of the double-stranded GFP fragment. The heat map displays relative fluorescence intensity after 3 hours normalized to no activator control in figure 2k in the main text. Error bars represent ± SEM, where n = 6 replicates. The experiments were repeated at least twice with n = 3 per experiment.

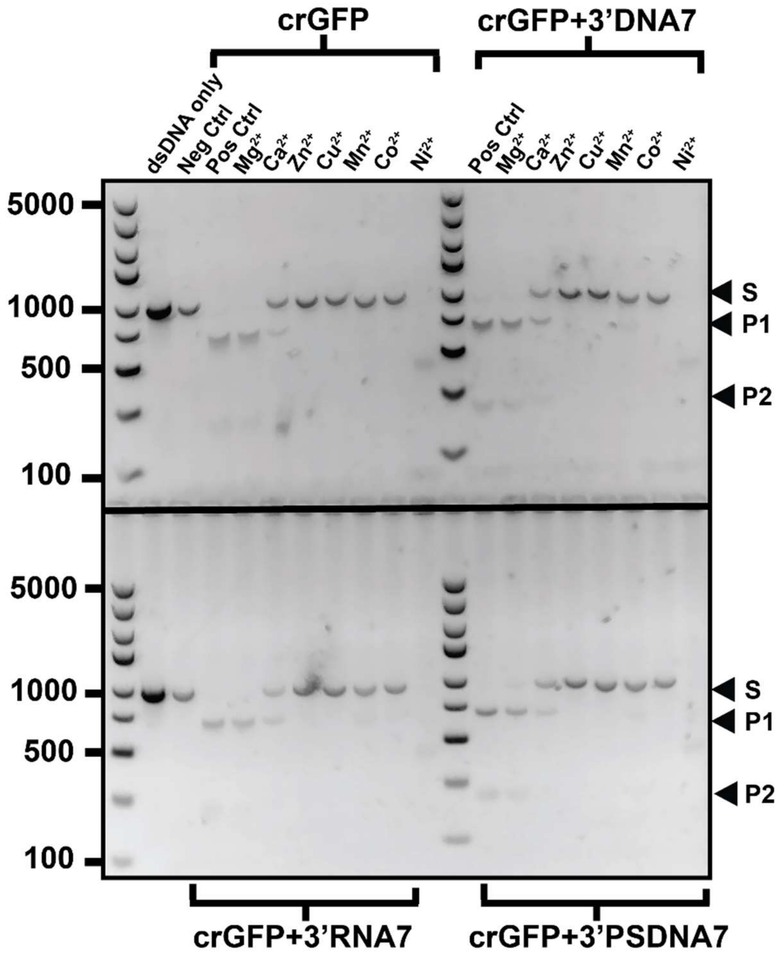

**Supplemental Figure 14.** The effect of a divalent ion on the cis-cleavage assay of LbCas12a with different crRNAs including wild-type crGFP (top-left quadrant), crGFP+3’DNA7 (top-right quadrant), crGFP+3’RNA7 (bottom-left quadrant), and crGFP+3’PSDNA7 (bottom-right quadrant). Neg Ctrl represents the negative control where crCon was used in the reaction mixture. Pos Ctrl represents the positive control where NEBuffer 2.1 was used. In this experiment, 100 nM of LbCpf1, 100 nM of crRNA, and 6.6 nM of DNA activator fragment were used.

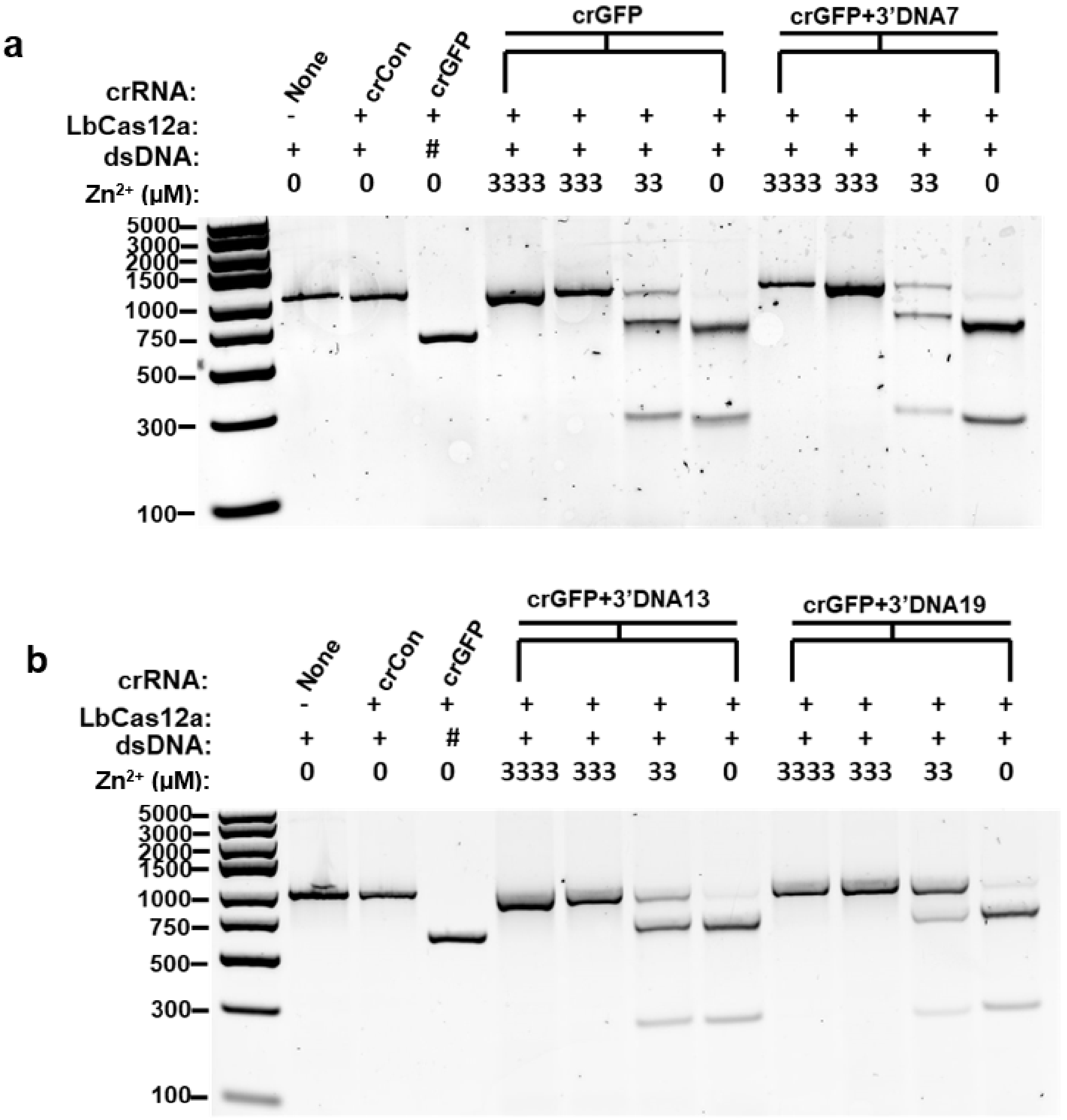

**Supplementary Figure 15.** The effect of $Zn^{2+}$ on Cis-cleavage assay of LbCas12a with different crRNAs in the presence of $Mg^{2+}$. 200 nM of LbCpf1, 200 nM of crRNA, 6.6 nM of DNA activator fragment, and 3mM of $Mg^{2+}$ were used. # stands for non-target dsDNA fragment. (**a**) crGFP and crGFP+3'DNA7. (**b**) crGFP+3'DNA13 and crGFP+3'DNA19.

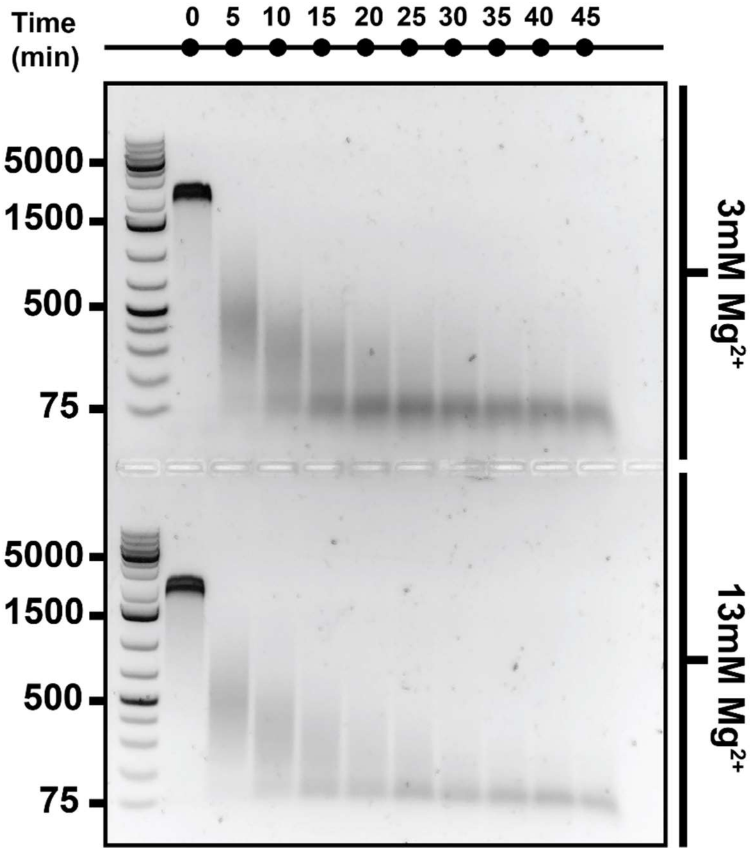

**Supplementary Figure 16.** Time-dependent cis-cleavage of LbCas12a on GFP in the presence of nonspecific ssDNA M13mp18 at varying $Mg^{2+}$ concentration. The reaction mixture was taken out every five minutes and quenched with 6X SDS-containing loading dye.

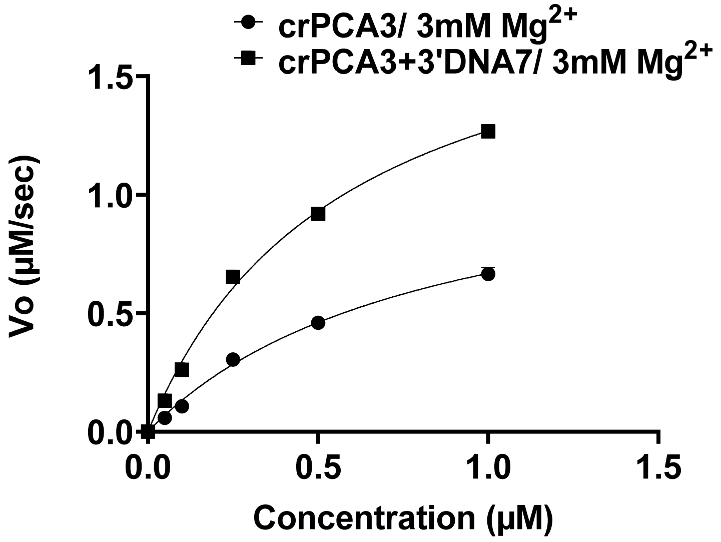

**Supplementary Figure 17.** Michaelis-Menten kinetic study of the wild type crPCA3 vs. crPCA3+3'DNA7. The graph shows initial velocity as a function of substrate concentration. In this case, the substrate used was /5HEX/TTA TT/3IABkFQ/.

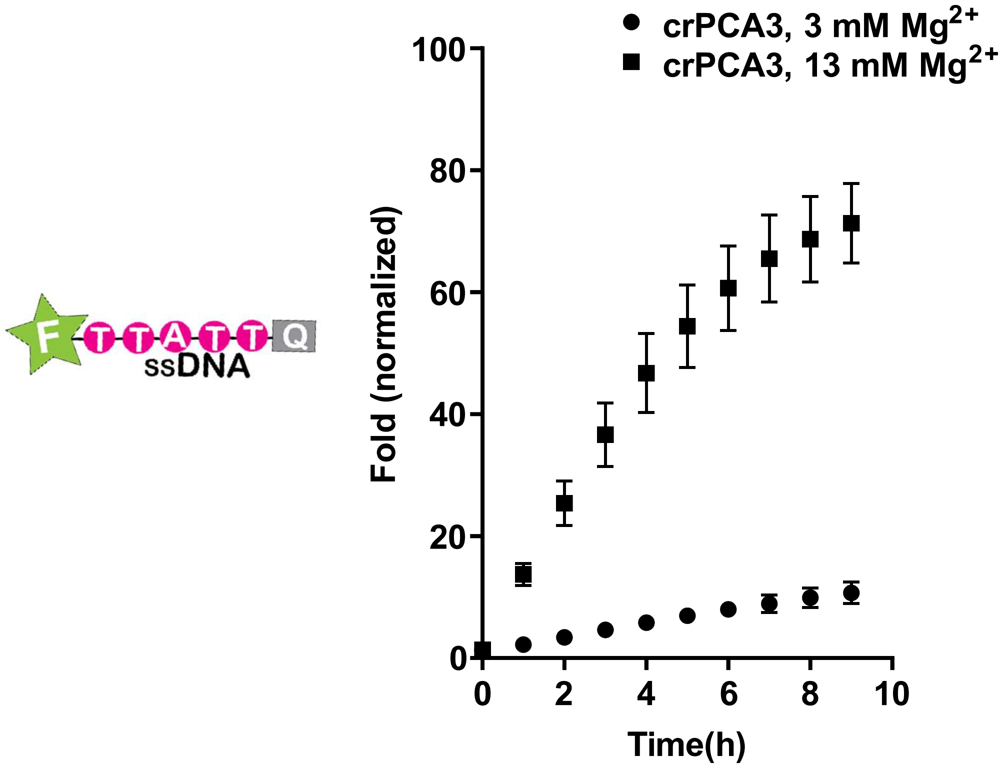

**Supplementary Figure 18.** Trans-cleavage activity of LbCas12 with modified 3' crPCA3 via fluorescence-quencher reporter assay FAM-TA at varying $Mg^{2+}$ concentration. FAM-TA is a reporter shown above. F stands for fluorophore, and Q stands for quencher. The fold in fluorescence was normalized by taking the ratio of background-corrected fluorescence signals of the samples with the activator to the corresponding samples without the activator. Error bars represent ± SEM, where n = 6 replicates. The experiments were repeated at least twice with n = 3 per experiment.

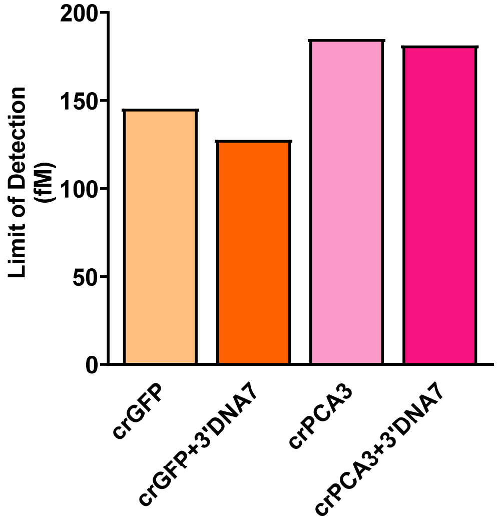

**Supplementary Figure 19. Limit of detection using modified crRNA/LbCas12a system.** Limit of detection in femtomolar at 13 mM $Mg^{2+}$ concentration. The reaction was carried out by adding simulated human urine spiked with either dsDNA GFP or PCA3 fragments to the LbCas12a reaction mixture.

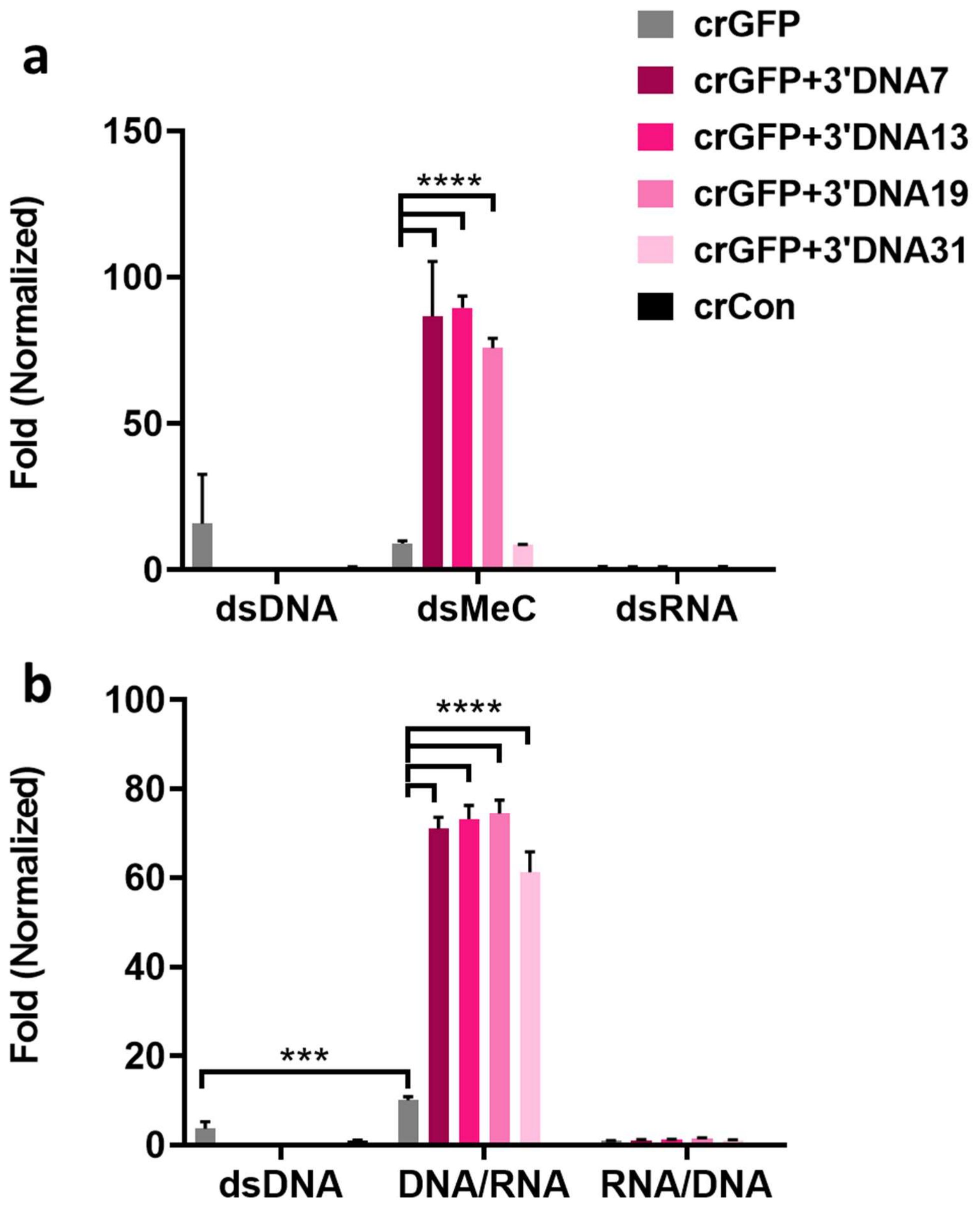

**Supplementary Figure 20.** LbCas12a trans-cleavage with different modified crRNAs and double-stranded activators (target and non-target strands annealed in the ratio of 1:1) measured by a fluorescence-quencher reporter assay (FAM-TA) in triplicates (n=3). The data shows results at 81 minutes for (A) dsDNA, dsMeC, sRNA, and (B) dsDNA, DNA/RNA, RNA/DNA. The values were normalized to their respective crRNA without activators. A two-way ANOVA test was used to calculate the p values where ****$p<0.0001$, ***$p<0.001$, and the error bars denote ±SD.

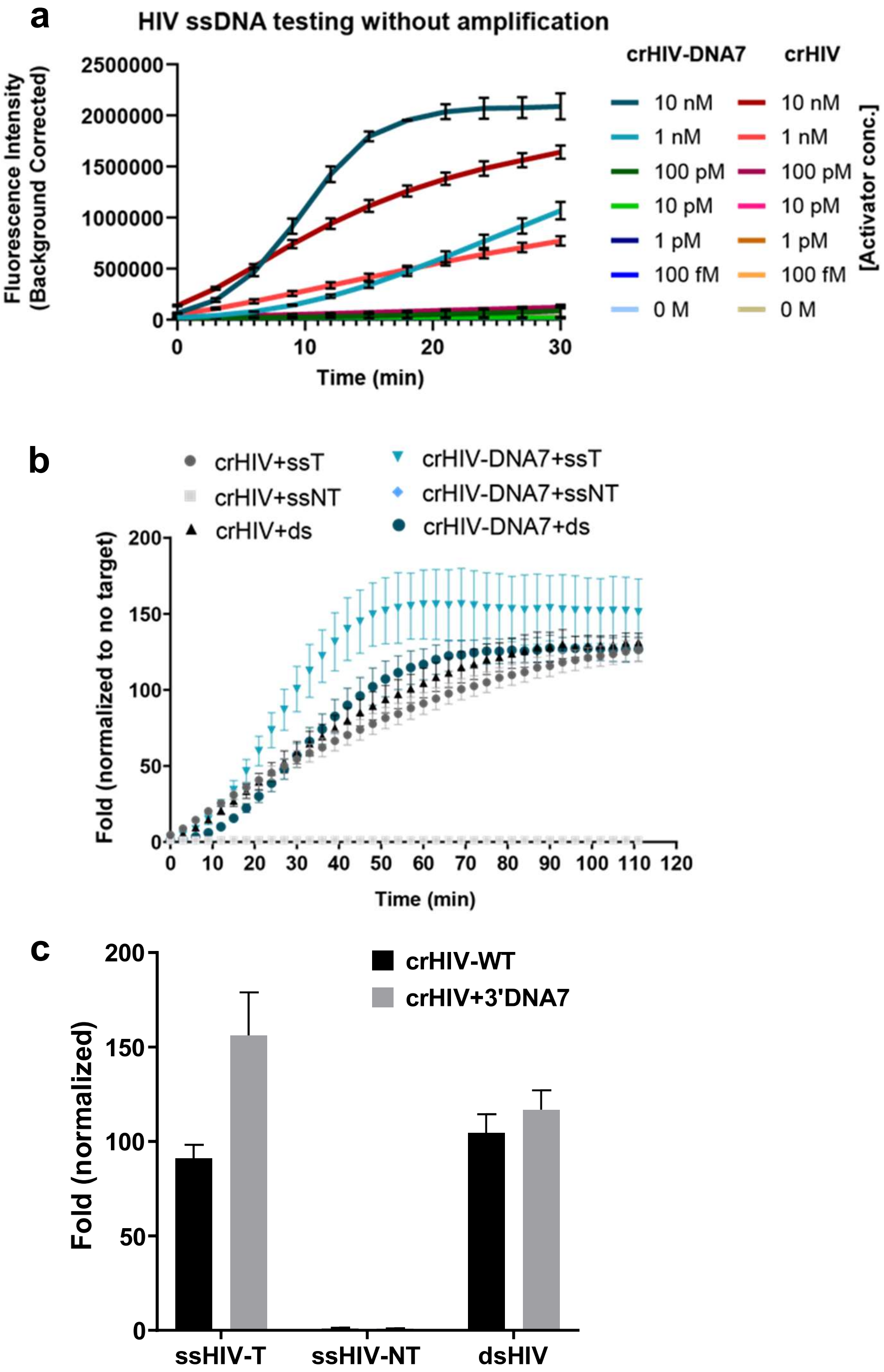

**Supplemental Figure 21.** (**a**) Trans-cleavage activity of regular crRNA vs. 3'DNA7 modified crRNA for detecting HIV ssDNA with LbCas12a over 30 minutes in the presence of various concentrations of HIV target dsDNA. Blank subtracted raw fluorescence intensities are indicated. (**b**) Fold change in fluorescence intensity in the presence vs. absence of target is indicated for various HIV ssDNA vs. dsDNA activators, where ssT indicates ssDNA target strand, ssNT indicates ssDNA non-target strand and ds indicates double-stranded DNA target, Mean ± SD (n=3). (**c**) Trans-cleavage of ssHIV using engineered CRISPR/LbCas12a. Comparison of single-stranded (ss) vs. double-stranded DNA (ds) targets analysis after 30 minutes is shown in bar graphs. HIV-1 ssDNA from Tat gene (IDT Technologies). The modified crRNA showed much higher sensitivity than the regular crRNA with fM detection limits within 30 minutes. Mean ± SEM (n = 3).

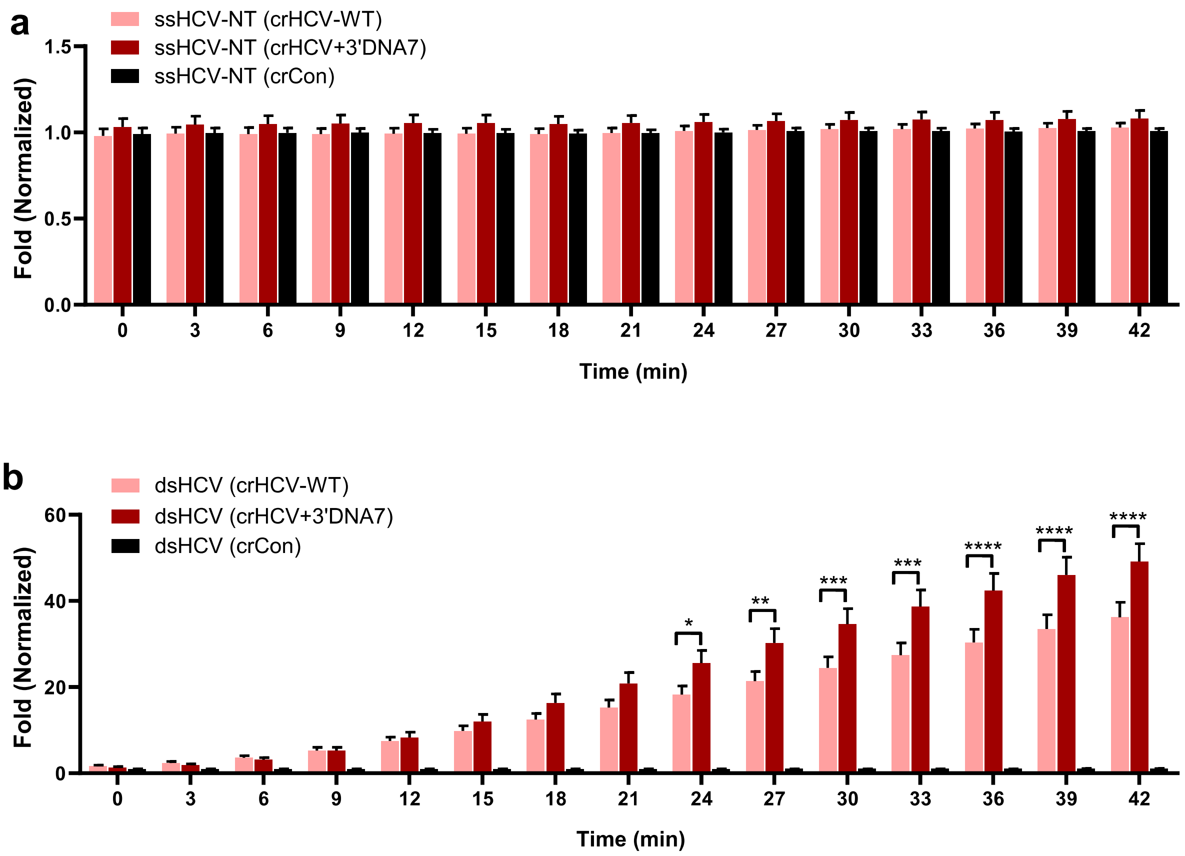

**Supplemental Figure 22.** Trans-cleavage activity of LbCas12a over time in the presence of 10 nM (100 pmols) of HCV non-target ssDNA (top) and HCV dsDNA (bottom). Using engineered crRNA with optimized CRISPR assay, detection of HCV target ssDNA was found to be 29 amols (290 fM, 100 □L) at 30 min, without target amplification, Mean ± SE, ANOVA (n=3, N=2, $^{ns}P > 0.05$, $^{*}P < 0.05$, $^{**}P < 0.01$, $^{***}P < 0.001$, $^{****}P < 0.0001$).

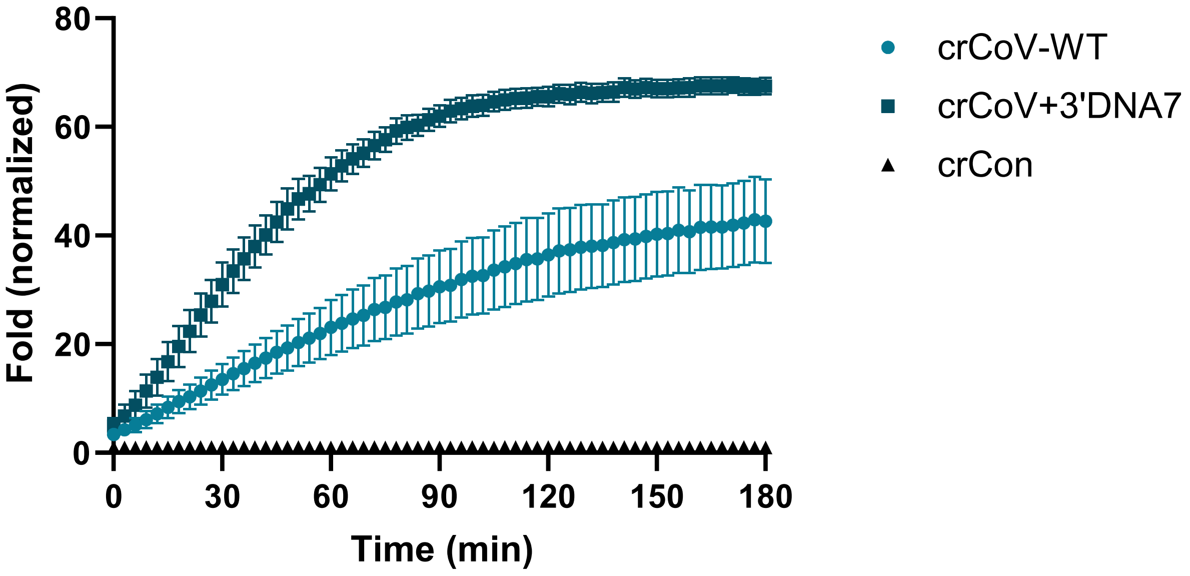

**Supplemental Figure 23.** Trans-cleavage activity of LbCas12a over time in the presence of 10 nM (100 pmols) of SARS-CoV-2 dsDNA target (bottom). Using engineered crRNA, containing 3'DNA7, and optimized CRISPR assay, the detection of SARS-COV-2 target ssDNA was found to be significantly faster compared to the wild type crRNA, without target amplification, Mean ± SE.

a

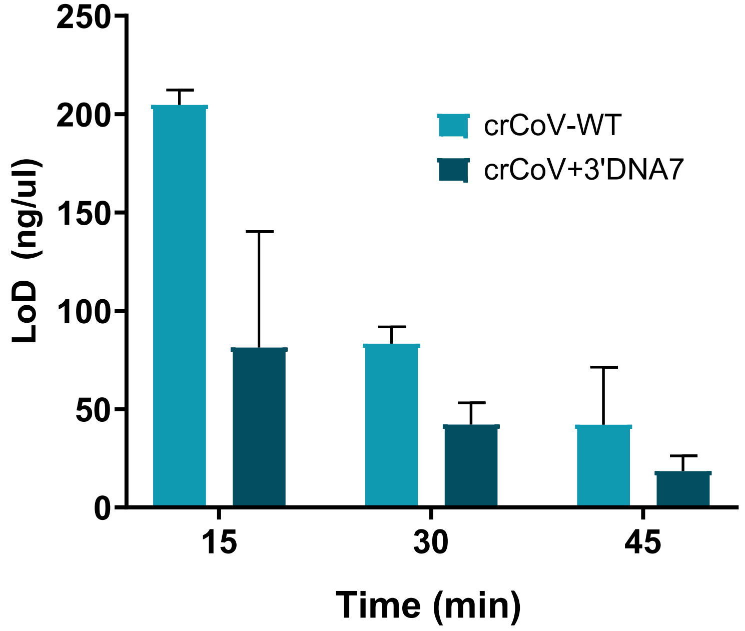

b

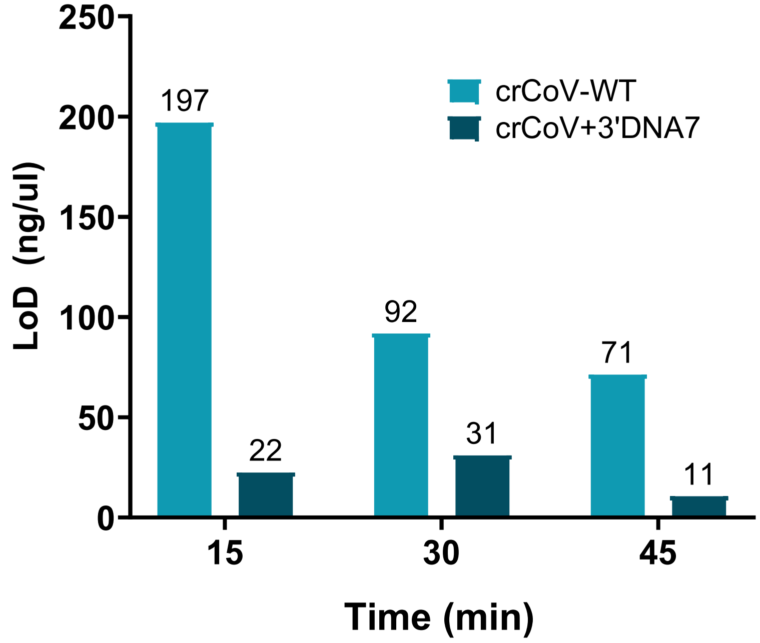

**Supplemental Figure 24.** Determination of lowest limit of detection for SARS-CoV-2 RNA using LbCas12a. To detect RNA, first a reverse transcriptase step was formed to convert RNA into DNA/RNA heteroduplex. The heteroduplex was detected by using an optimized CRISPR assay over time using either wild type crRNA (crCoV-WT) or engineered crRNA containing 3’DNA7 modifications (crCoV+3’DNA7). The limit of detection for SARS-COV-2 target RNA was found to be significantly lower with crCoV+3’DNA7, compared to the wild type crRNA, without target amplification. Top panel indicates Mean ± SE from two separate experiments (N=2). The bottom plot indicates a representative data with the lowest limit of detection.

a

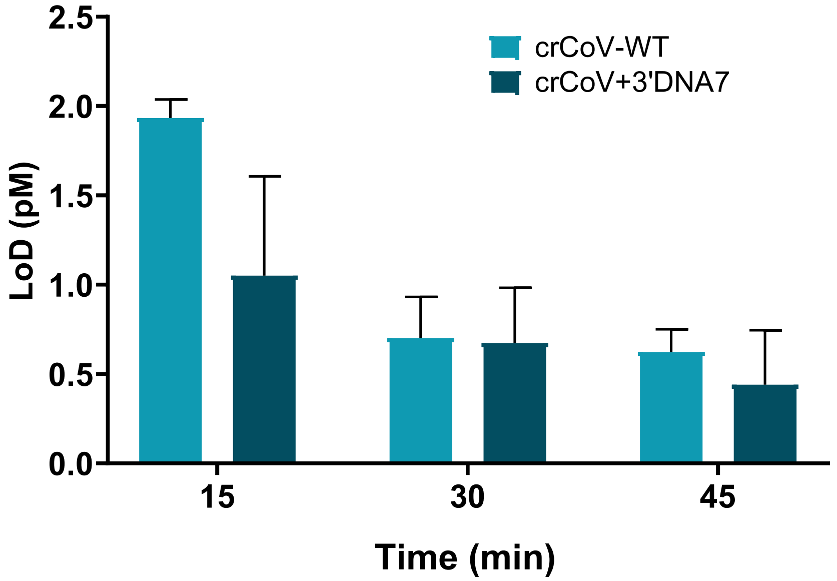

b

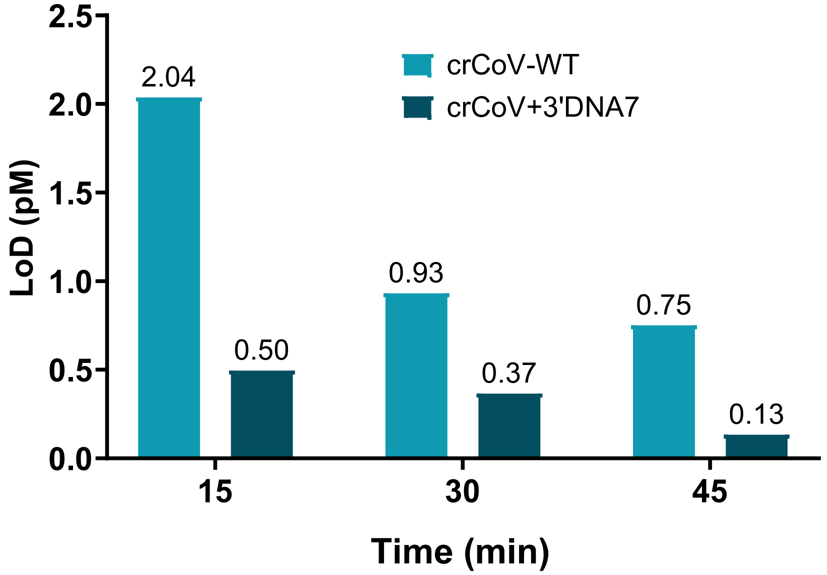

**Supplemental Figure 25.** Determination of lowest limit of detection for SARS-COV-2 dsDNA using LbCas12a with an optimized CRISPR assay over time using either wild type crRNA (crCoV-WT) or engineered crRNA containing 3'DNA7 modifications (crCoV+3'DNA7). The limit of detection for SARS-COV-2 target RNA was found to be significantly lower with crCoV+3'DNA7 (130 fM), compared to the crCoV-WT (750 fM) within 45 minutes, without target amplification. Top panel indicates Mean ± SE from two separate experiments (N=2). The bottom plot indicates a representative data with the lowest limit of detection.

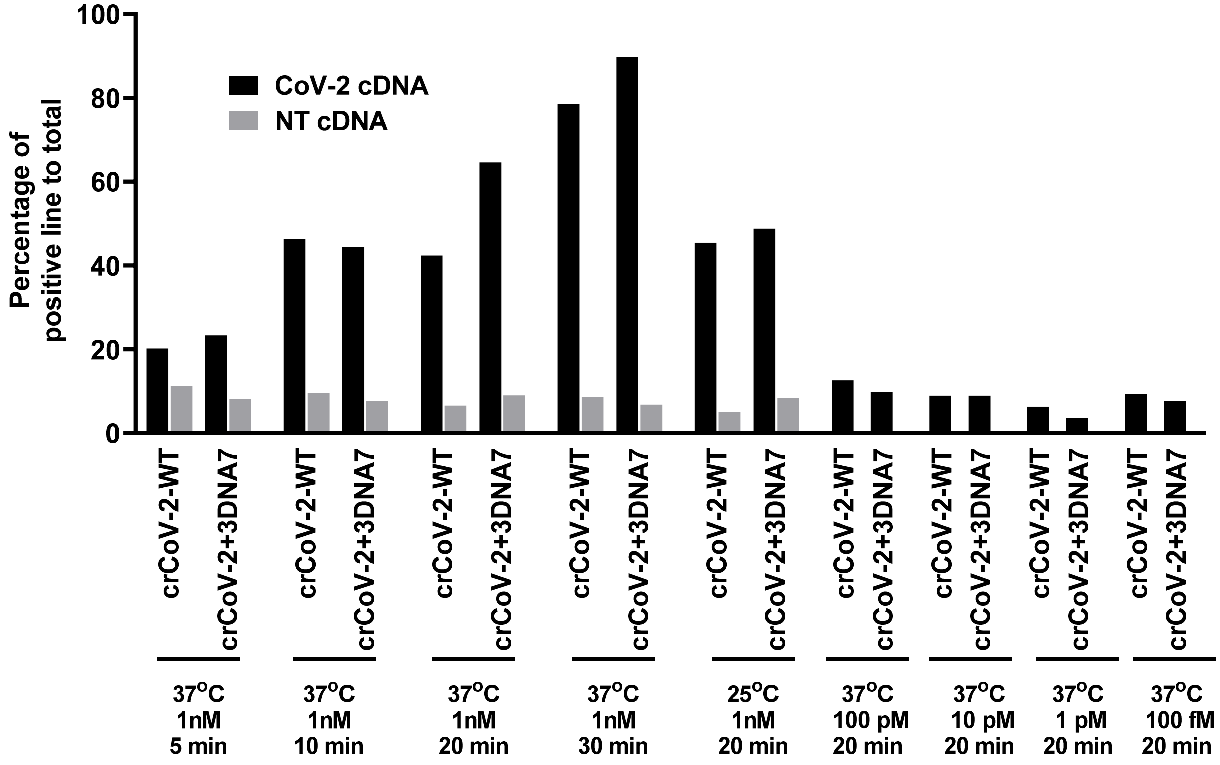

**Supplemental Figure 26.** Band intensity analysis of paper-strip test of LbCas12a targeting SARS-CoV-2. In this experiment, incubation time and temperature were varied (see materials and methods for more details). The paper-strips were scanned under Typhoon Amersham (GE healthcare) and analyzed using ImageJ.

a

b

c

d

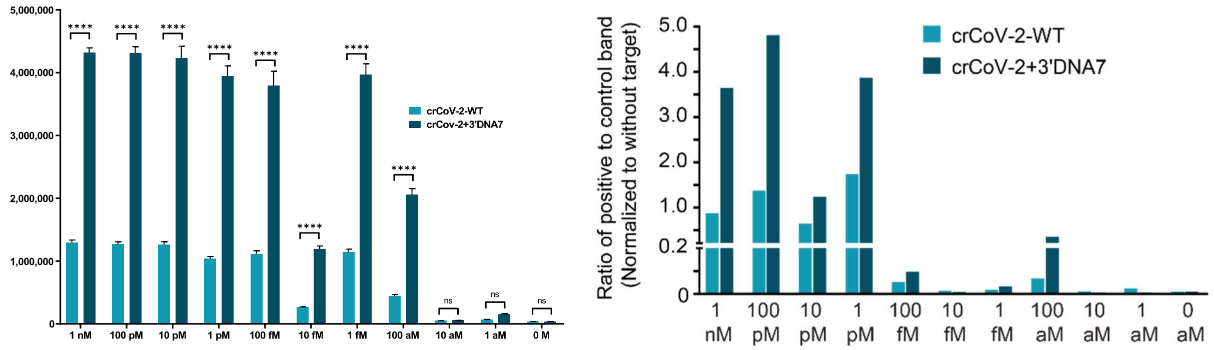

**Supplemental Figure 27.** Trans-cleavage activity of LbCas12a targeting SARS-CoV-2 with a RT-LAMP preamplification step via fluorescence-quencher reporter assay with FAM-TA**.** The data shown above are raw fluorescence signal measured by the fluorescence microplate reader Biotek Synergy 2. (a) and (b) are kinetics data in 15 min of crCoV-2-WT and crCoV-2+3'DNA7. In this experiment, the concentration of FQ reporter was doubled compared to previous experiments (100 nM). (c) is a single-point fluorescence signal extracted from (a) and (b). (d) Lateral flow assay with RT-LAMP preamplification step. The paper strips were scanned and analyzed using imageJ. Error bars represent ± SEM, where n = 6 replicates; two-way ANOVA test. The experiments were repeated at least twice with n = 3 per experiment.

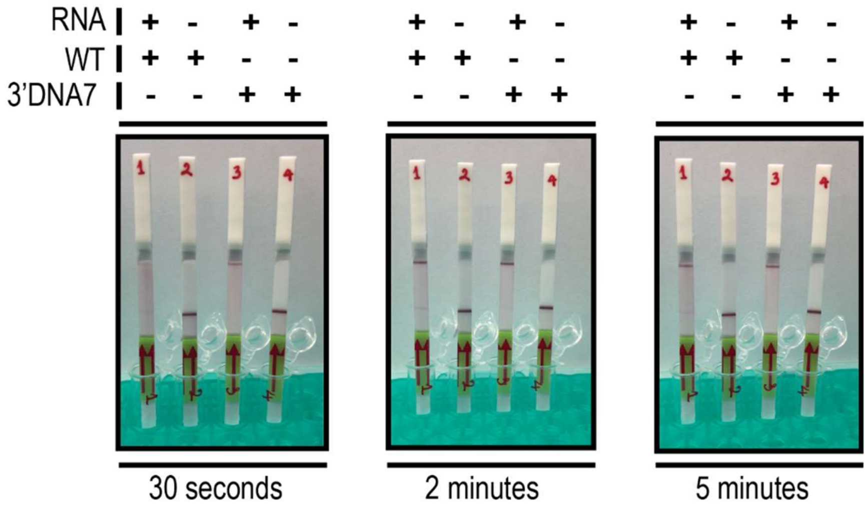

**Supplemental Figure 28.** A representation of time lapse pictures of the lateral flow assay targeting SARS-CoV-2 RNA N gene. The paper strips were immediately dipped into the LbCas12a reaction after the incubation. The testing concentration was 1 nM SARS-CoV-2 RNA.

**CRISPR RNAs (crRNAs)**

Selected crRNAs for AsCas12a

| Sequence Name | Sequence |
|---|---|
| AscrCon | /AITR1/rUrArA rUrUrU rCrUrA rCrUrC rUrUrG rUrArG rArUrC rGrUrU rArArU rCrGrC rGrUrA rUrArA rUrArC rGrG/AITR2/ |
| AscrGFP1 | /AltR1/rUrArA rUrUrU rCrUrA rCrUrC rUrUrG rUrArG rArUrC rGrUrC rGrCrC rGrUrC rCrArG rCrUrC rGrArC rC/AltR2/ |
| AscrGFP2-WT | /AltR1/rUrArA rUrUrU rCrUrA rCrUrC rUrUrG rUrArG rArUrC rUrCrA rGrGrG rCrGrG rArCrU rGrGrG rUrGrC rU/AltR2/ |
| AscrGFP2-WT-no-Alt | rUrArA rUrUrU rCrUrA rCrUrC rUrUrG rUrArG rArUrC rUrCrA rGrGrG rCrGrG rArCrU rGrGrG rUrGrC rU |
| AscrGFP2+3'DNA7 | rUrArA rUrUrU rCrUrA rCrUrC rUrUrG rUrArG rArUrC rUrCrA rGrGrG rCrGrG rArCrU rGrGrG rUrGrC rUTA TTA TT |

Selected crRNAs for FnCas12a

| | |
|---|---|
| FncrGFP2-WT | rUrArA rUrUrU rCrUrA rCrUrG rUrUrG rUrArG rArUrC rUrCrA rGrGrG rCrGrG rArCrU rGrGrG rUrGrC rU |
| FncrGFP2+3'DNA7 | rUrArA rUrUrU rCrUrA rCrUrG rUrUrG rUrArG rArUrC rUrCrA rGrGrG rCrGrG rArCrU rGrGrG rUrGrC rUTA TTA TT |
| FncrPCA3 | rUrArA rUrUrU rCrUrA rCrUrG rUrUrG rUrArG rArUrU rCrArC rCrCrC rUrGrC rCrArU rUrGrA rGrArU rG |

Selected crRNAs for LbCas12a

| | |
|---|---|
| LbcrCon (Neg-Ctrl-LbcrRNA) | rUrArA rUrUrU rCrUrA rCrUrA rArGrU rGrUrA rGrArU rCrGrU rUrArA rUrCrG rCrGrU rArUrA rArUrA rCrGrG |
| LbcrGFP1 | rUrArA rUrUrU rCrUrA rCrUrA rArGrU rGrUrA rGrArU rCrGrU rCrGrC rCrGrU rCrCrA rGrCrU rCrGrA rCrC |
| LbcrGFP2-WT | rUrArA rUrUrU rCrUrA rCrUrA rArGrU rGrUrA rGrArU rCrUrC rArGrG rGrCrG rGrArC rUrGrG rGrUrG rCrU |
| LbcrPCA3-WT | rUrArA rUrUrU rCrUrA rCrUrA rArGrU rGrUrA rGrArU rUrCrA rCrCrC rCrUrG rCrCrA rUrUrG rArGrA rUrG |
| LbcrPCA3+3'DNA7 | rUrArA rUrUrU rCrUrA rCrUrA rArGrU rGrUrA rGrArU rUrCrA rCrCrC rCrUrG rCrCrA rUrUrG rArGrA rUrGT ATT ATT |
| LbcrHIV-WT | rUrArArUrUrUrCrUrArCrUrArArGrUrGrUrArGrArUrCrCrUrUrGrGrUrGrG rGrUrGrCrUrArCrUrCrCrU |
| LbcrHIV+3'DNA7 | rUrArArUrUrUrCrUrArCrUrArArGrUrGrUrArGrArUrCrCrUrUrGrGrUrGrG rGrUrGrCrUrArCrUrCrCrUTATTATT |
| LbcrHCV-WT | rUrArArUrUrUrCrUrArCrUrArArGrUrGrUrArGrArUrUrGrCrUrCrArUrGrA rUrGrCrArCrGrGrUrCrUrA |
| LbcrHCV+3'DNA7 | rUrArArUrUrUrCrUrArCrUrArArGrUrGrUrArGrArUrUrGrCrUrCrArUrGr ArUrGrCrArCrGrGrUrCrUrATATTATT |
| crCoV-2-WT | rUrArArUrUrUrCrUrArCrUrArArGrUrGrUrArGrArUrGrUrGrGrArCrCrCr UrCrArGrArUrUrCrArArCrU |
| crCoV-2+3’DNA7 | rUrArArUrUrUrCrUrArCrUrArArGrUrGrUrArGrArUrGrUrGrGrArCrCrCr UrCrArGrArUrUrCrArArCrUTATTATT |

3’ DNA modified crGFP2 for LbCas12a

| Sequence Name | Sequence |
|---|---|
| LbcrGFP2+3’DNA7 | rUrArA rUrUrU rCrUrA rCrUrA rArGrU rGrUrA rGrArU rCrUrC rArGrG rGrCrG rGrArC rUrGrG rGrUrG rCrUT ATT ATT |
| LbcrGFP2+3’DNA13 | rUrArA rUrUrU rCrUrA rCrUrA rArGrU rGrUrA rGrArU rCrUrC rArGrG rGrCrG rGrArC rUrGrG rGrUrG rCrUT ATT ATT ATT ATT |
| LbcrGFP2+3’DNA19 | rUrArA rUrUrU rCrUrA rCrUrA rArGrU rGrUrA rGrArU rCrUrC rArGrG rGrCrG rGrArC rUrGrG rGrUrG rCrUT ATT ATT ATT ATT ATT ATT |
| LbcrGFP2+3’DNA31 | rUrArA rUrUrU rCrUrA rCrUrA rArGrU rGrUrA rGrArU rCrUrC rArGrG rGrCrG rGrArC rUrGrG rGrUrG rCrUT ATT ATT ATT ATT ATT ATT ATT ATT ATT ATT |
| LbcrGFP2+3’DNA7(GC) | rUrArA rUrUrU rCrUrA rCrUrA rArGrU rGrUrA rGrArU rCrUrC rArGrG rGrCrG rGrArC rUrGrG rGrUrG rCrUC GCC GCC |

5’ DNA modified crGFP2 for LbCas12a

| Sequence Name | Sequence |
|---|---|
| LbcrGFP2+5'DNA7 | TTA TTA TrUrA rArUrU rUrCrU rArCrU rArArG rUrGrU rArGrA rUrCrU rCrArG rGrGrC rGrGrA rCrUrG rGrGrU rGrCrU |
| Tru-LbcrGFP2+5'DNA7 | TTA TTA TrArA rUrUrU rCrUrA rCrUrA rArGrU rGrUrA rGrArU rCrUrC rArGrG rGrCrG rGrArC rUrGrG rGrUrG rCrU |
| LbcrGFP2+5'DNA13 | TTA TTA TTA TTA TrUrA rArUrU rUrCrU rArCrU rArArG rUrGrU rArGrA rUrCrU rCrArG rGrGrC rGrGrA rCrUrG rGrGrU rGrCrU |
| LbcrGFP2+5'DNA19 | TTA TTA TTA TTA TTA TTA TrUrA rArUrU rUrCrU rArCrU rArArG rUrGrU rArGrA rUrCrU rCrArG rGrGrC rGrGrA rCrUrG rGrGrU rGrCrU |
| LbcrGFP2+5'DNA31 | TTA TTA TTA TTA TTA TTA TTA TTA TTA TTA TrUrA rArUrU rUrCrU rArCrU rArArG rUrGrU rArGrA rUrCrU rCrArG rGrGrC rGrGrA rCrUrG rGrGrU rGrCrU |

3’ PSDNA modified crGFP2 for LbCas12a

| Sequence Name | Sequence |
|---|---|
| LbcrGFP2+3'PSDNA7 | rUrArA rUrUrU rCrUrA rCrUrA rArGrU rGrUrA rGrArU rCrUrC rArGrG rGrCrG rGrArC rUrGrG rGrUrG rCrU*T* A*T*T* A*T*T |
| LbcrGFP2+3'PSDNA13 | rUrArA rUrUrU rCrUrA rCrUrA rArGrU rGrUrA rGrArU rCrUrC rArGrG rGrCrG rGrArC rUrGrG rGrUrG rCrU*T* A*T*T* A*T*T* A*T*T* A*T*T |
| LbcrGFP2+3'PSDNA19 | rUrArA rUrUrU rCrUrA rCrUrA rArGrU rGrUrA rGrArU rCrUrC rArGrG rGrCrG rGrArC rUrGrG rGrUrG rCrU*T* A*T*T* A*T*T* A*T*T* A*T*T* A*T*T* A*T*T |

5’ PSDNA modified crGFP2 for LbCas12a

| Sequence Name | Sequence |
|---|---|
| LbcrGFP2+5'PSDNA7 | T*T*A* T*T*A* T*rUrA rArUrU rUrCrU rArCrU rArArG rUrGrU rArGrA rUrCrU rCrArG rGrGrC rGrGrA rCrUrG rGrGrU rGrCrU |
| LbcrGFP2+5'PSDNA13 | T*T*A* T*T*A* T*T*A* T*T*A* T*rUrA rArUrU rUrCrU rArCrU rArArG rUrGrU rArGrA rUrCrU rCrArG rGrGrC rGrGrA rCrUrG rGrGrU rGrCrU |

| | |
|---|---|
| LbcrGFP2+5'PSDNA19 | T*T*A* T*T*A* T*T*A* T*T*A* T*T*A* T*T*A* T*rUrA rArUrU rUrCrU rArCrU rArArG rUrGrU rArGrA rUrCrU rCrArG rGrGrC rGrGrA rCrUrG rGrGrU rGrCrU |

3' RNA modified crGFP2 for LbCas12a

| **Sequence Name** | **Sequence** |
|---|---|
| LbcrGFP2+3'RNA7 | rUrArA rUrUrU rCrUrA rCrUrA rArGrU rGrUrA rGrArU rCrUrC rArGrG rGrCrG rGrArC rUrGrG rGrUrG rCrUrU rArUrU rArUrU |
| LbcrGFP2+3'RNA13 | rUrArA rUrUrU rCrUrA rCrUrA rArGrU rGrUrA rGrArU rCrUrC rArGrG rGrCrG rGrArC rUrGrG rGrUrG rCrUrU rArUrU rArUrU rArUrU rArUrU |
| LbcrGFP2+3'RNA19 | rUrArA rUrUrU rCrUrA rCrUrA rArGrU rGrUrA rGrArU rCrUrC rArGrG rGrCrG rGrArC rUrGrG rGrUrG rCrUrU rArUrU rArUrU rArUrU rArUrU rArUrU rArUrU |

5' RNA modified crGFP2 for LbCas12a

| **Sequence Name** | **Sequence** |
|---|---|
| LbcrGFP2+5'RNA7 | rUrUrA rUrUrA rUrUrA rArUrU rUrCrU rArCrU rArArG rUrGrU rArGrA rUrCrU rCrArG rGrGrC rGrGrA rCrUrG rGrGrU rGrCrU |
| LbcrGFP2+5'RNA13 | rUrUrA rUrUrA rUrUrA rUrUrA rUrUrA rArUrU rUrCrU rArCrU rArArG rUrGrU rArGrA rUrCrU rCrArG rGrGrC rGrGrA rCrUrG rGrGrU rGrCrU |
| LbcrGFP2+5'RNA19 | rUrUrA rUrUrA rUrUrA rUrUrA rUrUrA rUrUrA rUrUrA rArUrU rUrCrU rArCrU rArArG rUrGrU rArGrA rUrCrU rCrArG rGrGrC rGrGrA rCrUrG rGrGrU rGrCrU |

**Activator DNA&RNA**

| Sequence Name | Sequence |
|---|---|
| DD3/PCA3-40nt-T-ssDNA | AGA CTA CAG ACA TCT CAA TGG CAG GGG TGA GAA ATA AGA A |
| DD3/PCA3-40nt-T-ssDNA – complement | TTC TTA TTT CTC ACC CCT GCC ATT GAG ATG TCT GTA GTC T |
| TTATT sequence-13mer | TATTATTATTATT |
| DD3-PCA3-gene-250bp-transcript variant 1 | CAA GAT AAA TAA GTG AAG AGC TAG TCC GCT GTG AGT CTC CTC AGT GAC ACA GGG CTG GAT CAC CAT CGA CGG CAC TTT CTG AGT ACT CAG TGC AGC AAA GAA AGA CTA CAG ACA TCT CAA TGG CAG GGG TGA GAA ATA AGA AAG GCT GCT GAC TTT ACC ATC TGA GGC CAC ACA TCT GCT GAA ATG GAG ATA ATT AAC ATC ACT AGA AAC AGC AAG ATG ACA ATA TAA TGT CTA AGT AGT GAC ATG TTT T |
| LbCas12a-Activator-HIV1-RNA | rUrUrCrUrCrUrCrUrGrCrArCrCrArCrUrCrUrUrCrUrCrUrUrUrGrCrCrUrUrGrGrUrGrGrGrUrGrCrUrArCrUrCrCrUrArArUrGrGrUrUrCrArArUrUrUrU |
| LbCas12a-HIV-DNA-T-Activator | AAAATTGAACCATTAGGAGTAGCACCCACCAAGGCAAAGAGAAGAGTGGTGCAGAGAGAA |
| LbCas12a-HIV-DNA-NT-Activator | TTCTCTCTGCACCACTCTTCTCTTTGCCTTGGTGGGTGCTACTCCTAATGGTTCAATTTT |
| HCV_RNA_reference_Genome | rGrCrCrUrUrGrUrGrGrUrArCrUrGrCrCrUrGrArUrArGrGrGrUrGrCrUrUrGrCrGrArGrUrGrCrCrCrCrGrGrGrArGrGrUrCrUrCrGrUrArGrArCrCrGrUrGrCrArUrCrArUrGrArGrCrArCrArArArUrCrCrUrArArArCrCrUrC |
| HCV_DNA_act_T | CCTCTAATACGACTCACTATAGGCGTTGGGTTGCGAACGGCCTTGTGGTACTGCCTGATAGGGTGCTTGCGAGTGCCCCGGGAGGTCTCGTAGACCGTGCATCATGAGCACAAATCCTAAACCTC |
| HCV_DNA_act_NT | GAGGTTTAGGATTTGTGCTCATGATGCACGGTCTACGAGACCTCCCGGGGCACTCGCAAGCACCCTATCAGGCAGTACCACAAGGCCGTTCGCAACCCAACGCCTATAGTGAGTCGTATTAGAGG |
| 2019-nCoV_N_Positive Control | Plasmid CAT_10006625_2019-nCoV_N_Positive Control from IDT |

**Activator Primers**

| Sequence Name | Sequence |
|---|---|
| GFP-Act-NT-MedC | GGG GTC TTT G/iMe-dC/T /iMe-dC/AG GG/iMe-dC/ GGA /iMe-dC/TG GGT G/iMe-dC/T CAG GTA GTG G |
| GFP-Act-T-MedC | CCA CTA CCT GAG /iMe-dC/A/iMe-dC/ /iMe-dC//iMe-dC/A GT/iMe-dC/ /iMe-dC/G/iMe-dC/ /iMe-dC//iMe-dC/T GAG CAA AGA CCC C |
| GFP-40nt-T-heteroDNA-RNA | CCA CTA CCT GrArG rCrArC rCrCrA rGrUrC rCrGrC rCrCrU rGrArG CAA AGA CCC C |
| GFP-40nt-NT-heteroDNA-RNA | GGG GTC TTT GrCrU rCrArG rGrGrC rGrGrA rCrUrG rGrGrU rGrCrU CAG GTA GTG G |
| HIV_RNA_Primer_RT-Superscript | AAA ATT GAA CCA TTA GGA GTA GC |
| HIV_RNA_Primer_RT-MMLV | AAA ATT GAA CCA TTA GG |
| HIV-RNA-RP | TTC TCT CTG CAC CAC TCT TC |
| HCV_RP_gblock_also_RT1_LN | GAG GTT TAG GAT TTG TGC TCA T |
| HCV_RP_gblock_also_RT2_LN | GAG GTT TAG GAT TTG TGC TCA TGA |
| HCV_FP_1 | CCT CTA ATA CGA CTC ACT A |
| HCV_FP_2 | CCT CTA ATA CGA CTC A |
| HCV_PCR_cDNA_FP1 | GCG TTG GGT TGC GAA CGG CC |
| HCV_PCR_cDNA_FP2 | GCG TTG GGT TGC GAA CGG |
| HIV_RT_cDNA_FP | GCC TTG TGG TAC TGC CTG AT |
| 2019-nCoV_N1_FP | GAC CCC AAA ATC AGC GAA AT |
| 2019-nCoV_N1_RP | TCT GGT TAC TGC CAG TTG AAT CTG |
| 2019-nCoV_N1_T7FP | CCT CTA ATA CGA CTC ACT ATA GGA CCC CAA AAT CAG CGA AAT |
| 2019-nCoV_N1_T7RP | CCT CTA ATA CGA CTC ACT ATA GGT CTG GTT ACT GCC AGT TGA ATC TG |
| 2019-nCoV-N3_FP_LN | GGG AGC CTT GAA TAC ACC AAA A |
| 2019-nCoV_N3_RP_LN | TGT AGC ACG ATT GCA GCA TTG |
| 2019-nCoV_N2_FP_LN | TTA CAA ACA TTG GCC GCA AA |
| 2019-nCoV_N2_RP_LN | GCG CGA CAT TCC GAA GAA |
| F3_LAMP_CoV | AAC ACA AGC TTT CGG CAG |
| B3_LAMP_CoV | GCCTTGTCCTCGAGGGAAT |
| FIP_LAMP_CoV | CCACTGCGTTCTCCATTCTGGTAAATGCACCCCGCATTACG |
| BIP_ LAMP_CoV | CGCGATCAAAACAACGTCGGCCCTTGCCATGTTGAGTGAGA |

**FQ Substrates and Labeled crRNAs**

| Sequence Name | Sequence |
|---|---|
| ssDNA-FQ reporter1 | /56-FAM/TTA TT/3IABkFQ/ |
| Oligo 2 FAM-Biotin | /56-FAM/TTA TT/3Bio/ |
| 5'Cy5-3'RQ-FQ-Reporter | TAT TA/iCy5/T TAT T/3IAbRQSp/ |
| FQreporter-Hex-IowaFQ | /5HEX/TTA TT/3IABkFQ/ |
| ssDNA-FAM-FQ reporter1 | /56-FAM/TTA TT/3IABkFQ/ |
| FAM-GC-richFQ-Reporter | /56-FAM/CCG CC/3IABkFQ/ |
| FQreporter-FAM-ssRNA(rN)-IABFQ | /56-FAM/rCrCrG rCrC/3IABkFQ/ |
| FQ-reporter-FAM-ssRNA(UArich) | /56-FAM/rUrUrA rUrU/3IABkFQ/ |
| LbcrGFP2+3'DNA7-FAM | rUrArA rUrUrU rCrUrA rCrUrA rArGrU rGrUrA rGrArU rCrUrC rArGrG rGrCrG rGrArC rUrGrG rGrUrG rCrUT ATT ATT/36-FAM/ |
| 5'FAM-LbcrGFP2 | /56-FAM/rUrArA rUrUrU rCrUrA rCrUrA rArGrU rGrUrA rGrArU rCrUrC rArGrG rGrCrG rGrArC rUrGrG rGrUrG rCrU |
| LbcrGFP2-3'FAM | rUrArA rUrUrU rCrUrA rCrUrA rArGrU rGrUrA rGrArU rCrUrC rArGrG rGrCrG rGrArC rUrGrG rGrUrG rCrU/36-FAM/ |
| 5'FAM-LbcrGFP2+3'DNA13 | /56-FAM/rUrArA rUrUrU rCrUrA rCrUrA rArGrU rGrUrA rGrArU rCrUrC rArGrG rGrCrG rGrArC rUrGrG rGrUrG rCrUT ATT ATT ATT ATT |
| LbcrGFP2+3'DNA13-FAM | rUrArA rUrUrU rCrUrA rCrUrA rArGrU rGrUrA rGrArU rCrUrC rArGrG rGrCrG rGrArC rUrGrG rGrUrG rCrUT ATT ATT ATT ATT/36-FAM/ |
| 5'HEX-DNA19+LbcrGFP2 | /5HEX/TTA TTA TTA TTA TTA TTA TrUrA rArUrU rUrCrU rArCrU rArArG rUrGrU rArGrA rUrCrU rCrArG rGrGrC rGrGrA rCrUrG rGrGrU rGrCrU |
| LbcrGFP2+DNA-5'FQ-Cy5 (crGFP+5'DNA13+Cy5+DNA6+Iowa Black RQ) | /5IAbRQ/TTA TT/iCy5/A TTA TTA TTA TTA TrUrA rArUrU rUrCrU rArCrU rArArG rUrGrU rArGrA rUrCrU rCrArG rGrGrC rGrGrA rCrUrG rGrGrU rGrCrU |
| LbcrGFP2+DNA-3'FQ-Cy5 (crGFP+3'DNA7+Cy5+DNA6+Iowa Black RQ) | rUrArA rUrUrU rCrUrA rCrUrA rArGrU rGrUrA rGrArU rCrUrC rArGrG rGrCrG rGrArC rUrGrG rGrUrG rCrUT ATT ATT A/iCy5/TT ATT /3IAbRQSp/ |

**Primers**

| Sequence Name | Sequence |
|---|---|
| LbcrGFP2-3'DNA7-Primer-15 | AAT AAT AAG CAC CCA GTC CGC C |
| LbcrGFP2-3'DNA7-Primer-14 | AAT AAT AAG CAC CCA GTC CGC |
| LbcrGFP2-3'DNA7-Primer-13 | AAT AAT AAG CAC CCA GTC CG |
| LbcrGFP2-3'DNA7-Primer-12 | AAT AAT AAG CAC CCA GTC C |
| LbcrGFP2-3'DNA7-Primer-11 | AAT AAT AAG CAC CCA GTC |
| LbcrGFP2-3'DNA7-Primer-10 | AAT AAT AAG CAC CCA GT |
| LbcrGFP2-3'DNA7-Primer-9 | AAT AAT AAG CAC CCA G |
| LbcrGFP2-3'DNA7-Primer-8 | AAT AAT AAG CAC CCA |
| LbcrGFP2-3'DNA7-Primer-7 | AAT AAT AAG CAC CC |
| LbcrGFP2-3'DNA7-Primer-6 | AAT AAT AAG CAC C |
| LbcrGFP2-3'DNA7-Primer-5 | AAT AAT AAG CAC |
| LbcrGFP2-3'DNA7-Primer-4 | AAT AAT AAG CA |
| LbcrGFP2-Primer-15 | AGC ACC CAG TCC GCC |
| LbcrGFP2-Primer-14 | AGC ACC CAG TCC GC |
| LbcrGFP2-Primer-13 | AGC ACC CAG TCC G |
| LbcrGFP2-Primer-12 | AGC ACC CAG TCC |
| LbcrGFP2-Primer-11 | AGC ACC CAG TC |
| LbcrGFP2-Primer-10 | AGC ACC CAG T |
| LbcrGFP2-Primer-9 | AGC ACC CAG |
| LbcrGFP2-Primer-8 | AGC ACC CA |
| RPA-PCA3-FP1 | AGT ACT CAG TGC AGC AAA GAA AGA CTA CAG |
| RPA-PCA3-RP1 | ACA TTA TAT TGT CAT CTT GCT GTT TCT AGT GAT |
| RPA-PCA3-FP2 | AGT GAA GAG CTA GTC CGC TGT GAG TCT CCT |
| RPA-PCA3-RP2 | CTG TTT CTA GTG ATG TTA ATT ATC TCC ATT TC |
| RPA-PCA3-FP3 | AAG AGC TAG TCC GCT GTG AGT CTC CTC AGT |
| RPA-PCA3-RP3 | GTT TCT AGT GAT GTT AAT TAT CTC CAT TTC AG |
| T7-Foward-primer1-RNA | CCT CTA ATA CGA CTC ACT ATA GGA ACG GCA TCA AGG TGA ACT |
| T7-Foward-primer2-RNA | CCT CTA ATA CGA CTC ACT ATA GGC GAC CAC TAC CAG CAG AAC A |
| Primer-EGFP-F490 | ACT TCA AGA TCC GCC ACA AC |
| Primer-EGFP-F473 | GAA CGG CAT CAA GGT GAA CT |
| Primer-EGFP-F536 | CGA CCA CTA CCA GCA GAA CA |

**Activators**

| Sequence Name | Sequence |
|---|---|
| Act-GFP-10nt-T-10nt | CCA CTA CCT GAG CAC CCA GTC CGC CCT GAG CAA AGA CCC C |
| Act-GFP-10nt-NT-10nt | GGG GTC TTT GCT CAG GGC GGA CTG GGT GCT CAG GTA GTG G |
| Act-GFP-10nt-NT-5MeC-10nt | GGG GTC TTT G/iMe-dC/T CAG GGC GGA CTG GGT GCT CAG GTA GTG G |
| Act-GFP-10nt-T-5MeC-10nt | CCA CTA CCT GAG CAC CCA GTC CGC CCT GAG /iMe-dC/AA AGA CCC C |
| Act-GFP-mut-1 | CCA CTA CCT GGG CAC CCA GTC CGC CCT GAG CAA AGA CCC C |
| Act-GFP-mut-2 | CCA CTA CCT GAA CAC CCA GTC CGC CCT GAG CAA AGA CCC C |
| Act-GFP-mut-3 | CCA CTA CCT GAG TAC CCA GTC CGC CCT GAG CAA AGA CCC C |
| Act-GFP-mut-4 | CCA CTA CCT GAG CGC CCA GTC CGC CCT GAG CAA AGA CCC C |
| Act-GFP-mut-5 | CCA CTA CCT GAG CAT CCA GTC CGC CCT GAG CAA AGA CCC C |
| Act-GFP-mut-6 | CCA CTA CCT GAG CAC TCA GTC CGC CCT GAG CAA AGA CCC C |
| Act-GFP-mut-7 | CCA CTA CCT GAG CAC CTA GTC CGC CCT GAG CAA AGA CCC C |
| Act-GFP-mut-8 | CCA CTA CCT GAG CAC CCG GTC CGC CCT GAG CAA AGA CCC C |
| Act-GFP-mut-9 | CCA CTA CCT GAG CAC CCA ATC CGC CCT GAG CAA AGA CCC C |
| Act-GFP-mut-10 | CCA CTA CCT GAG CAC CCA GCC CGC CCT GAG CAA AGA CCC C |
| Act-GFP-mut-11 | CCA CTA CCT GAG CAC CCA GTT CGC CCT GAG CAA AGA CCC C |
| Act-GFP-mut-12 | CCA CTA CCT GAG CAC CCA GTC TGC CCT GAG CAA AGA CCC C |
| Act-GFP-mut-13 | CCA CTA CCT GAG CAC CCA GTC CAC CCT GAG CAA AGA CCC C |
| Act-GFP-mut-14 | CCA CTA CCT GAG CAC CCA GTC CGT CCT GAG CAA AGA CCC C |
| Act-GFP-mut-15 | CCA CTA CCT GAG CAC CCA GTC CGC TCT GAG CAA AGA CCC C |
| Act-GFP-mut-16 | CCA CTA CCT GAG CAC CCA GTC CGC CTT GAG CAA AGA CCC C |
| Act-GFP-mut-17 | CCA CTA CCT GAG CAC CCA GTC CGC CCC GAG CAA AGA CCC C |
| Act-GFP-mut-18 | CCA CTA CCT GAG CAC CCA GTC CGC CCT AAG CAA AGA CCC C |
| Act-GFP-mut-19 | CCA CTA CCT GAG CAC CCA GTC CGC CCT GGG CAA AGA CCC C |

| | |
|---|---|
| Act-GFP-mut-20 | CCA CTA CCT GAG CAC CCA GTC CGC CCT GAA CAA AGA CCC C |
| Act-GFP-mut-1 - complement | GGG GTC TTT GCT CAG GGC GGA CTG GGT GCC CAG GTA GTG G |
| Act-GFP-mut-2 – complement | GGG GTC TTT GCT CAG GGC GGA CTG GGT GTT CAG GTA GTG G |
| Act-GFP-mut-3 – complement | GGG GTC TTT GCT CAG GGC GGA CTG GGT ACT CAG GTA GTG G |
| Act-GFP-mut-4 – complement | GGG GTC TTT GCT CAG GGC GGA CTG GGC GCT CAG GTA GTG G |
| Act-GFP-mut-5 - complement | GGG GTC TTT GCT CAG GGC GGA CTG GAT GCT CAG GTA GTG G |
| Act-GFP-mut-6 - complement | GGG GTC TTT GCT CAG GGC GGA CTG AGT GCT CAG GTA GTG G |
| Act-GFP-mut-7 - complement | GGG GTC TTT GCT CAG GGC GGA CTA GGT GCT CAG GTA GTG G |
| Act-GFP-mut-8 - complement | GGG GTC TTT GCT CAG GGC GGA CCG GGT GCT CAG GTA GTG G |
| Act-GFP-mut-9 – complement | GGG GTC TTT GCT CAG GGC GGA TTG GGT GCT CAG GTA GTG G |
| Act-GFP-mut-10 – complement | GGG GTC TTT GCT CAG GGC GGG CTG GGT GCT CAG GTA GTG G |
| Act-GFP-mut-11 – complement | GGG GTC TTT GCT CAG GGC GAA CTG GGT GCT CAG GTA GTG G |
| Act-GFP-mut-12 – complement | GGG GTC TTT GCT CAG GGC AGA CTG GGT GCT CAG GTA GTG G |
| Act-GFP-mut-13 – complement | GGG GTC TTT GCT CAG GGT GGA CTG GGT GCT CAG GTA GTG G |
| Act-GFP-mut-14 – complement | GGG GTC TTT GCT CAG GAC GGA CTG GGT GCT CAG GTA GTG G |
| Act-GFP-mut-15 – complement | GGG GTC TTT GCT CAG AGC GGA CTG GGT GCT CAG GTA GTG G |
| Act-GFP-mut-16 – complement | GGG GTC TTT GCT CAA GGC GGA CTG GGT GCT CAG GTA GTG G |
| Act-GFP-mut-17 – complement | GGG GTC TTT GCT CGG GGC GGA CTG GGT GCT CAG GTA GTG G |
| Act-GFP-mut-18 – complement | GGG GTC TTT GCT TAG GGC GGA CTG GGT GCT CAG GTA GTG G |
| Act-GFP-mut-19 – complement | GGG GTC TTT GCC CAG GGC GGA CTG GGT GCT CAG GTA GTG G |
| Act-GFP-mut-20 - complement | GGG GTC TTT GTT CAG GGC GGA CTG GGT GCT CAG GTA GTG G |
